# Adolescent Brain Cognitive Development (ABCD) Community MRI Collection and Utilities

**DOI:** 10.1101/2021.07.09.451638

**Authors:** Eric Feczko, Greg Conan, Scott Marek, Brenden Tervo-Clemmens, Michaela Cordova, Olivia Doyle, Eric Earl, Anders Perrone, Darrick Sturgeon, Rachel Klein, Gareth Harman, Dakota Kilamovich, Robert Hermosillo, Oscar Miranda-Dominguez, Azeez Adebimpe, Maxwell Bertolero, Matthew Cieslak, Sydney Covitz, Timothy Hendrickson, Anthony C. Juliano, Kathy Snider, Lucille A. Moore, Johnny Uriartel, Alice M. Graham, Finn Calabro, Monica D. Rosenberg, Kristina M. Rapuano, BJ Casey, Richard Watts, Donald Hagler, Wesley K. Thompson, Thomas E. Nichols, Elizabeth Hoffman, Beatriz Luna, Hugh Garavan, Theodore D. Satterthwaite, Sarah Feldstein Ewing, Bonnie Nagel, Nico U.F. Dosenbach, Damien A. Fair

## Abstract

The Adolescent Brain Cognitive Development Study (ABCD), a 10 year longitudinal neuroimaging study of the largest population based and demographically distributed cohort of 9-10 year olds (N=11,877), was designed to overcome reproducibility limitations of prior child mental health studies. Besides the fantastic wealth of research opportunities, the extremely large size of the ABCD data set also creates enormous data storage, processing, and analysis challenges for researchers. To ensure data privacy and safety, researchers are not currently able to share neuroimaging data derivatives through the central repository at the National Data Archive (NDA). However, sharing derived data amongst researchers laterally can powerfully accelerate scientific progress, to ensure the maximum public benefit is derived from the ABCD study. To simultaneously promote collaboration and data safety, we developed the ABCD-BIDS Community Collection (ABCC), which includes both curated processed data and software utilities for further analyses. The ABCC also enables researchers to upload their own custom-processed versions of ABCD data and derivatives for sharing with the research community. This NeuroResource is meant to serve as the companion guide for the ABCC. In section we describe the ABCC. Section II highlights ABCC utilities that help researchers access, share, and analyze ABCD data, while section III provides two exemplar reproducibility analyses using ABCC utilities. We hope that adoption of the ABCC’s data-safe, open-science framework will boost access and reproducibility, thus facilitating progress in child and adolescent mental health research.

## Introduction

On September 30th, 2015 the National Institute of Health (NIH) and associated partners, such as the Centers for Disease Control (CDC) and the National Science Foundation (NSF), launched an unprecedented push to study adolescent brain health and development - the Adolescent Brain Cognitive Development study (ABCD). ABCD is the largest long-term study of brain development in the United States. It comprises 21 sites (Casey et al., 2018; Volkow et al., 2018) and has now enrolled 11,877 participants between 9-10 years of age. Each participant will be followed through young adulthood to determine how various experiences throughout adolescence affect numerous behavioral, academic, and health outcomes. Magnetic Resonance Imaging (MRI) is employed to monitor structural and functional brain development every 2 years. At the end of this 12-year effort, ~25,000,000 non-biological measurements will have been collected, and each participant will have had 5 MRI scanning sessions consisting of 12 distinct MR image acquisitions, totaling > 700,000 MRI scans. In contrast to the ABCD study, typical sample sizes in neuroimaging studies, according to a recent review, include less than 40 participants (Szucs and Ioannidis, 2020).

The ABCD‘s enormous data resources are being provided to the community in near real-time to maximize their utility and impact. However, the size and scope of the ABCD dataset poses a significant challenge for many interested investigators. The computing resources, processing time, data storage demands, complexity of nested variables (e.g., family members, study site), and statistical intricacies all can serve as barriers to entry for researchers conducting ABCD analyses. Added security needed to address privacy concerns further complicates accessibility.

### Section I: ABCD-BIDS Community MRI Collection (ABCC)

#### ABCD Data Access: Maximizing Privacy, Security and Accessibility

The ABCD study must acquire Protected Health Information (PHI). Such information can be used to link participant identification to collected measures, causing participant re-identification, so a large data set with PHI that has already received substantial media attention requires extremely strong privacy and security measures (Mazor et al., 2017; von Thenen et al., 2019). It is critical that all ABCD data are secured, de-identified and used appropriately. Therefore, the NIH, the ABCD Coordinating Center, and the ABCD Data Analysis, Informatics, & Resource Center (DAIRC) have put into place a secure framework that allows for sharing of raw data and derived data types between the ABCD study team and the community via the National Data Archive (NDA) (Auchter et al., 2018).

The ABCD team releases demographic, behavioral and raw neuroimaging data in near ‘real-time’ in the form of Fast Track Data Releases (Auchter et al., 2018). In addition the ABCD’s DAIRC provides a yearly updated condensed ‘tabulated’ data release, which includes neuroimaging measures derived using the DAIRC-maintained processing pipeline, as well as a ‘minimally pre-processed’ dataset where distortion correction, bias field correction, and motion correction have been applied (Hagler et al., 2019)(ABCD Data Collection 2573). The DAIRC also maintains tools such as the Data Exploration and Analysis Portal (DEAP) (Bartsch et al., 2014) to provide an analytical framework that facilitates standardized practices for statistical analyses.

Critically important ABCD data security measures included in the data use agreement (https://nda.nih.gov/ndapublicweb/Documents/NDA+Data+Access+Request+DUC+FINAL.pdf) provide investigators access to the data, but prevent them from sharing any individual-specific derived measures outside of the NDA. Thus, the only mechanism to share subject specific ABCD-derived data is through the NDA. These necessary restrictions on lateral data sharing greatly limit the risks of irresponsible data usage, such as participant re-identification, but limit also collaboration within the research community. The process of organizing, maintaining, and uploading newly derived ABCD measures to the NDA can be computationally intensive.

For these reasons, we created an additional ‘community share’ for the easier distribution of ABCD-derived data (Feczko et al., 2020a). The *ABCD BIDS Community Collection* (ABCC, ABCD-3165) enables community contributions and usage, with standardized formatting (BIDS derivatives structure), a governance structure to maintain compliance with the NDA’s important data usage standards (Mazor et al., 2017; von Thenen et al., 2019), and analytic utilities to improve data accessibility and ease of use.

The ABCC data (ABCD-3165) and ABCC analytic utilities were designed to 1) supplement current data sharing via collection 3165 with an accessible platform for community sharing of derived data via our NDA BIDS prepare and upload tool(Earl and Fair, 2021) 2) assist investigators in conducting state-of-the art statistical analyses using software packages designed to work with ABCD-3165, and 3) provide integrated data and analytical utilities for result verification.

In this NeuroResource, we detail how the original ABCC data (ABCD-3165) were generated (Section I), introduce helpful utilities (Section II) designed for use with the ABCC (3165), and demonstrate how these utilities can be applied (Section III) to the ABCC data (ABCD-3165). Specifically, we: 1) Describe the ABCD-BIDS processing pipeline(Earl et al., 2020a), built on the principles of the Human Connectome Project (HCP) pipeline (Glasser et al., 2013; Marek et al., 2019), used to generate the ABCC (ABCD-3165). 2) Detail the ABCC (ABCD-3165) data structure with input and processed derivative data in a standardized format (BIDS: Brain Imaging Data Structure) for easy accessibility and analysis. 3) We highlight the ABCD Reproducible Matched Samples(Cordova et al., 2020a) (ARMS) designed for accurate, demographics-matched, replicability testing in ABCD. 4) Deliver a software package that performs vertex and voxelwise Brain-Wide Association studies (BWAS) and controls for batch effects and nested variables (Guillaume et al., 2014) called the Sandwich Estimator for Neuroimaging Data (SEND) (Feczko et al., 2020b). 5) We conclude by discussing sample size considerations for measuring reproducibility and effect sizes using specific ABCC data examples.

#### ABCD-BIDS: Surface-based, HCP-style MRI Processing Pipeline

##### Brain Imaging Data Structure (BIDS) Format Increases Reproducibility by Standardizing Input Data

In neuroimaging, there has historically been little consensus on the formatting, storing, and naming of files. The lack of standards has made data sharing difficult and contributed to the creation of many, often redundant tools for analyzing and processing imaging data. The Brain Imaging Data Structure (BIDS) provides neuroimaging researchers with a standard for organizing and describing MRI datasets (Gorgolewski et al., 2016). The BIDS standard utilizes file formats widely compatible with existing software, unifies the majority of current best practices and captures the metadata necessary for most data processing operations (Gorgolewski et al., 2016). Through such data format standardization, BIDS generates an ecosystem for the improved integration of input data, derived data, and software applications (BIDS Applications [Apps] -- (Gorgolewski et al., 2017b)). The BIDS standardization allows for easier sharing and reuse, as well as greater pipeline reproducibility and analysis repeatability. The ABCD-BIDS pipeline (Earl et al., 2020a) is a BIDS application which allows researchers who adopt the BIDS standard to process their data with minimal user input while minimizing human error (Gorgolewski et al., 2017b).

Currently, BIDS input data that have passed Quality Assurance (QA) by the NDA (https://nda.nih.gov/about/standard-operating-procedures.html#sop5) for ABCD are provided through the ABCC (N=10,038). The success of this effort and community demand for BIDS input data via the ABCC, has now led to the inclusion of BIDS structured data for the Fast Track raw data releases beginning at the end of 2020. The ABCC will continue to curate BIDS input data that have passed initial QA.

##### The ABCD-BIDS Pipeline Was Designed for Maximum Versatility

The ABCD-BIDS (Earl et al., 2020a) processing pipeline (v.1.0) used to generate the ABCC(Earl et al., 2020b)(ABCD-3165) is an updated version of The Human Connectome Project (HCP - (Glasser et al., 2013)) MRI pipeline. The HCP MRI pipeline has been a successful platform and utilized across many studies. It comprises five stages (Figure 1; see supplemental for a detailed pipeline description). 1) PreFreesurfer, 2) Freesurfer, 3) PostFreesurfer, 4) Volume, and 5) Surface. ‘PreFreesurfer’ performs brain extraction, denoising, and normalizes the structural data to a standard template. ‘Freesurfer’ segments subcortical structures, reconstructs native surfaces from the normalized structural data, and registers the surfaces to the template surface. ‘PostFreesurfer’ converts native surfaces into HCP-compatible format (i.e. CIFTIs). ‘Volume’ registers the functional/diffusion MRI data to the volumetric standard template through the normalized structural data. ‘Surface’ projects the functional/diffusion data to the template-space surfaces.

**Figure 1.**
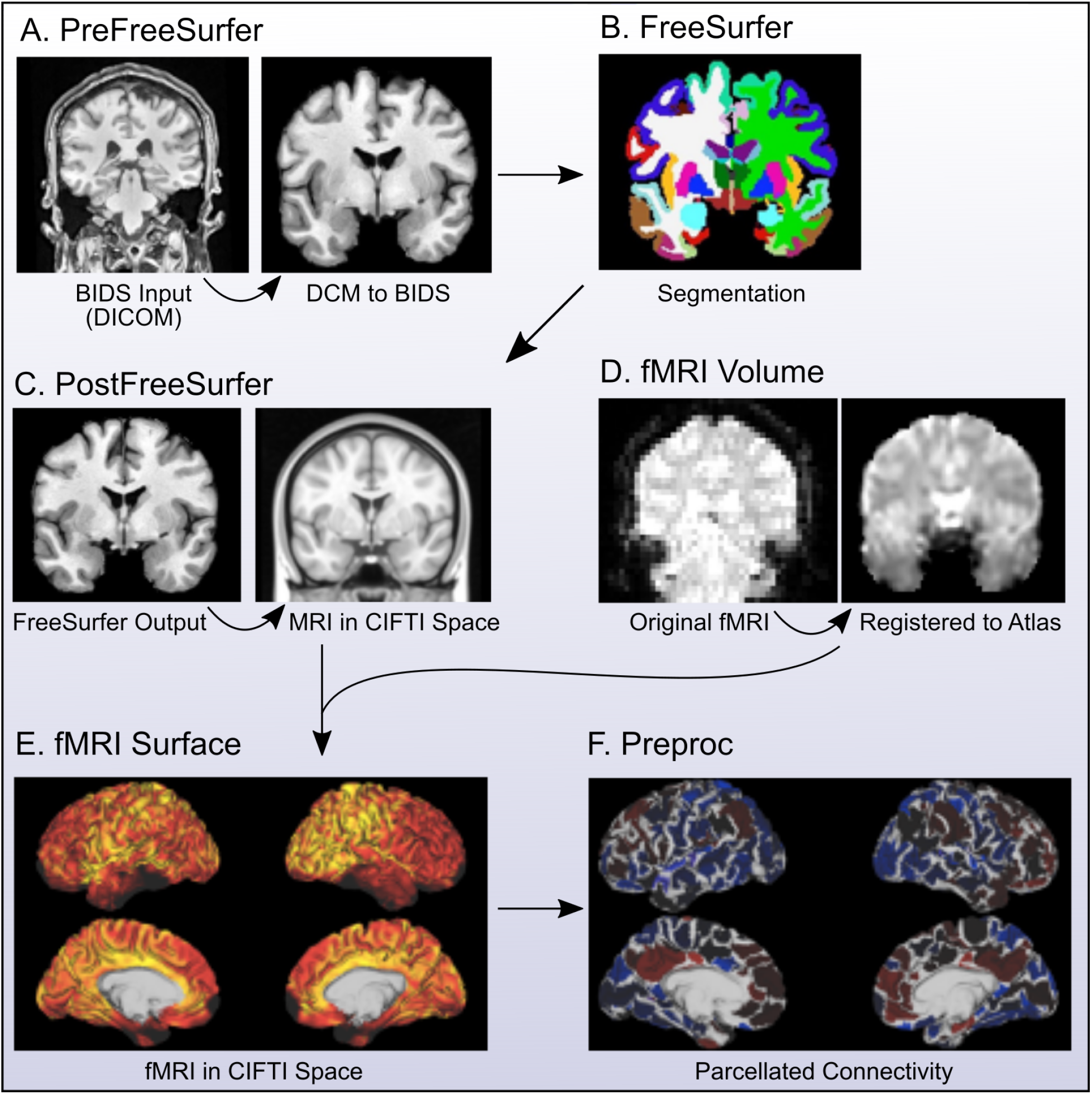
Workflow of the ABCD-BIDS Pipeline. The DCAN ABCD-BIDS pipeline comprises six primary stages: **(A) PreFreesurfer.** BIDS input anatomical data are normalized to an MNI template (“normalized” here refers collectively to masking, denoising, bias correction, and registration, see supplemental materials for a detailed list of steps). (**B) Freesurfer.** Normalized anatomical data are segmented and surfaces are generated via FreeSurfer. **(C) PostFreeSurfer.** Freesurfer and PreFreesurfer outputs are converted to a standard CIFTI template space. **(D) fMRIVolume.** BIDS formatted functional MRI data are registered to a standard MNI template **(E) fMRISurface.** Functional MRI data are then projected to the CIFTI standard template **(F) Preproc.** Functional MRI data undergo additional preprocessing beyond minimally preprocessed data. In addition, parcellated timeseries data are also generated.

Unfortunately, the HCP pipeline was primarily designed to comply with Siemens scanners and was optimized for the data collection procedures and hardware of the HCP study itself (Glasser et al., 2013). To attenuate scanner manufacturer effects and increase the usability of our processing pipeline across all MR platforms and data, we adopted and modified the HCP pipeline to accommodate GE, Phillips, and Siemens scanners and head coils from all 21 ABCD sites (Glasser et al., 2013; Marek et al., 2019). Our ABCD-BIDS pipeline automatically generates a series of outputs, including cortical surfaces, myelin maps, and processed functional MRI (fMRI) data. We embedded the ABCD-BIDS pipeline within a docker environment (Gorgolewski et al., 2017b) and as a BIDS App for ease of use, to attenuate computing environment effects, and enhance reproducibility (Glatard et al., 2015). With the docker-contained ABCD-BIDS pipeline, users can implement this pipeline immediately reducing concerns for limited reproducibility (Glatard et al., 2015).

An ABCD-BIDS pipeline workflow diagram (Figure 1) shows each step’s primary input and output. BIDS input anatomical data (i.e., T1 or both T1 and T2) serve as the primary input to PreFreesurfer (Figure 1A), which normalizes the anatomical data and rigidly aligns the data to a template, preserving the native structure. The normalized data are input to FreeSurfer, which produces and refines native cortical surfaces (Fischl, 2012) (Figure 1B). The processed anatomical data are converted to the CIFTI format in PostFreeSurfer (Figure 1C). fMRI data are brought into this CIFTI space by normalizing the volumetric data in fMRIVolume (Figure 1D) and projecting the normalized data onto the CIFTI format in fMRISurface (Figure 1E). DCAN BOLD preprocessing (DBP), or “Preproc”, uses the DCAN (Developmental Cognition and Neuroimaging) Labs connectivity preprocessing program (Miranda-Domínguez et al., 2020) on the fMRI CIFTI data, resulting in both dense (dtseries) and parcellated (ptseries) CIFTI datasets (Figure 1F).

The ABCD-BIDS pipeline differs from the HCP pipeline in PreFreesurfer, Freesurfer, PostFreesurfer, and Volume. These changes are detailed in supplementary material (see: “Additional modifications for ABCD BIDS pipeline”), however, the two most important updates are highlighted here. In PreFreesurfer, we modified the HCP pipeline to make the inclusion of T2w images optional. If a high quality T1w but low quality T2w were acquired, making the T2w image optional can allow for the inclusion of data sets that would otherwise have had to be excluded. We also perform the nonlinear registration to the standard atlas in PostFreeSurfer, which increases the effectiveness of the registration. This improvement is secondary to the refined brain mask generated from the Freesurfer outputs. Additionally, the ANTS deformable SyN algorithm performs the nonlinear registration instead of the FNIRT elastic deformation algorithm. A previous comparison of over 12 registration methods across multiple datasets show that ANTs consistently outperforms FNIRT on every performance metric (Ou et al., 2014). The ABCD-BIDS pipeline also provides the option to use a study-specific template as an intermediate image for atlas registration, which can help improve the registration in populations with systematic structural differences (e.g. aging populations with large ventricles).

In addition, two stages are added after “fMRISurface”. Step 6 - “Preproc” functional connectivity pre-processing (DCAN BOLD processing), based on Power *et. al.* (2014), utilizes our filtering process for the motion estimates, which separate true head motion from factitious motion secondary to magnetic field changes due to breathing (Fair et al., 2020). Step 7 - “Executive Summary”, summarizes standard quality control outputs. Our quality control summaries provide users a browser-interface for reviewing the processed data. These are described in detail in SI material. For more information please consult our online documentation (https://collection3165.readthedocs.io/en/stable/pipeline/). Efforts are ongoing to merge the functionality of the ABCD-BIDS pipelines with the popular fMRIprep utilities (Esteban et al., 2018, 2019a, 2019b), and XCP(Ciric et al., 2018) putting it in the NiPreps ecosystem(Esteban et al., 2019b) to maximize community input, standardization, and reproducibility. Future contributors and releases in the third quarter of 2021 will include derivatives based on fMRIprep version 20.0.7 (Esteban et al., 2019a, 2019b), and QSIPrep version 0.12.2 (Cieslak et al., 2020, 2021). In addition, task related contrasts are scheduled for a second quarter release date as well, based on the ABCD-FSLtask pipeline(Juliano et al., 2021) (see supplementary material: “ABCD task-fmri pipeline”).

#### The ABCD-BIDS Community Collection (ABCC; ABCD-3165) Simplifies Access to ABCD Community Derived Data

##### ABCC data (ABCD-3165) Utilizes the Standard BIDS Derivative Scheme

Similar to BIDS input data discussed above, ABCC (ABCD-3165) output data are also organized into a standardized “BIDS derivatives” format (Gorgolewski et al., 2016). BIDS derivatives are outputs of processing pipelines, such as the ABCD-BIDS pipeline, which capture data and metadata sufficient for a researcher to understand and reuse those outputs in subsequent processing. Across institutions different research groups are using distinct formatting standards when sharing neuroimaging data (Poldrack et al., 2017). As the amount of data and the breadth of sources continues to increase, organizing such data into a reproducible standard takes significant work. File naming and directory organization are inconsistent between sources thus making data manipulation and analysis more cumbersome. Metadata about processing routines are variable, often necessitating requests for clarification from the providers of the data. It is inefficient for many research groups to carry out this work in parallel. BIDS standardized derivatives ensure that processing outputs are organized and named consistently, along with standardized metadata. Therefore, data sharing under the BIDS structure provides a way for researchers to share processing pipeline outputs with full transparency as to how they were generated.

The BIDS standards continue to evolve and are implemented via a community driven governance and decision making structure (https://bids.neuroimaging.io/governance.html). New derivatives not already under the BIDS standard specifications go through the ‘BIDS extensions’ process (https://bids.neuroimaging.io/get_involved). Because converting processed data to BIDS derivatives is cumbersome when working with many MR sessions, we developed a stand-alone ABCC utility, called Filemapper, to reduce the burden of formatting data for BIDS-derivative schemes(Earl et al., 2020c). Filemapper maps any file/folder structure (e.g. HCP or BIDS outputs) into any other file/folder structure (e.g. BIDS or HCP outputs). Filemapper uses JavaScript Object Notation files (JSONs) to copy and rename files and folders from one location to another. If BIDS standards change, JSONs will be updated to reflect the changes. For ABCC data (ABCD-3165), Filemapper is a powerful generalized interface for mapping file structures between standards, such as converting the ABCD-BIDS pipeline outputs into BIDS format, consistent with the BIDS data dictionary. Standardizing processing inputs (BIDS), outputs (BIDS-derivatives) and processing software (BIDS Apps) greatly reduces the workload for researchers by making data and code interchangeable. BIDS standardization also facilitates the ease of data and code sharing and reduces the risk of human error, especially when collaborating with other research groups. All data shared through the ABCC and approved via our governance structure (see: “How to Contribute.”) will be in BIDS derivative formats and have passed BIDS validation (Figure 2).

**Figure 2.**
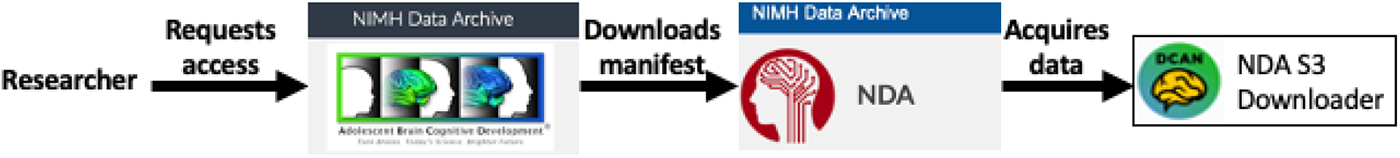
Workflow of ABCD-3165 (ABCC) request. Researchers first request access to the NDA’s ABCD collections via the NIMH data archive ABCD page. After getting NDA access to ABCD collections, including ABCD-3165, researchers can download the ABCD-3165 data manifest from the NDA. To acquire data subsets from the ABCD-3165, researchers can use the NDA S3 downloader(Earl et al., 2020b).

##### Accessing the ABCC Data

As mentioned previously, the ABCC is hosted by the NIH. Therefore, in order to obtain access to the ABCC data, prospective users will need to complete 5 steps (Figure 2). First, the prospective user must create a NDA account (https://nda.nih.gov/user/create_account.html). Users must be affiliated with an NIH-recognized institution with Federalwide Assurance for protection of human subjects and have a research specific request for the data. The user must submit and complete a data use certification (DUC) agreement for access to the NIMH data archive (https://nda.nih.gov/training/module?trainingModuleId=training.access&slideId=slide.access.intro). The DUC forbids “re-identification”; researchers cannot attempt to reveal the identity of any individual using the data provided. The signing official for the user’s affiliated institution must sign the DUC, along with the user. The user then submits the DUC to the NDA, who will review the requested approval. Once access is approved, the ABCC data can be downloaded from the NDA as specified below.

ABCC data (ABCD-3165) in BIDS format can currently be downloaded in two different ways. First, if one wants to access a specific subset of the data, we recommend downloading the “associated files” spreadsheet with Amazon Web Services (AWS) Simple Storage Service (S3) links, from NDA Collection 3165. Researchers can then download subject-specific folders using the GitHub nda-abcd-s3-downloader(Earl et al., 2020b).

The ABCC (ABCD-3165) does not contain the original, unprocessed DICOM files. To access the original DICOM files, one must download ABCD Fast Track Data from the NDA (ABCD 2573) and unpack them into BIDS using the ABCD-STUDY abcd-dicom2bids GitHub repository. The “Abcd dicom2bids”(Earl et al., 2020d) tool can pull DICOMs and externally collected data, such as behavior collected during task designed fMRI (e.g. E-Prime files) from the NDA’s “fast-track” data (ABCD 2573). It also unpacks, converts, and BIDS-standardizes the fast-track data so that they match ABCC (ABCD-3165).

##### ABCC Data Share: The “Big Data” Pediatric MRI Dataset

The ABCC data share is a population-based, demographically diverse, and stable MRI dataset comprising 11,877 children. The data provided are under version-control and stamped with release notes to ensure that analyses are reproducible. The available data contain multiple domains (e.g. raw BIDS formatted, processed, and derivative data) and are ready for subsequent processing or analysis steps (see: Sections II and III, below). Here we depict a subset for easy visualization (Table 1), a more complete description can be found in the supplemental materials (Table S1). The root folder (Table 1; yellow) describes the entire ABCD-BIDS (3165) dataset and repository contents. Derivatives (Table 1; orange) are separated by various pipeline outputs. For example, the ABCD-BIDS pipeline outputs are at the first level divided into functional (e.g. ####-task-rest_bold_desc-filtered-timeseries.dtseries.nii) and anatomical (e.g. ####_hemi-L_space-MNI_mesh-fsLR32k_midthickness.surf.gii) outputs. Functional outputs include quality control metrics such as frame-by-frame motion estimates, parcellated timeseries comprising different ROI sets, and atlas transformed dense timeseries files comprising over 91,000 grey-ordinates, and connectivity matrices. Anatomical outputs include cortical surface metrics and mid-thickness surface files in native and atlas space. For manual quality control, an html file in the ses-ID folder (####.html) can be accessed to view the executive summary in a web browser. The images folder contains the images viewed on the executive summary html file. External data produced during the scan, such as behavioral task performance data, can be found in the “sourcedata” folder (Table 1; blue). Finally, investigators interested in re-processing the data can access the BIDS inputs from each subject’s session folder (Table 1; green).

**Table 1.**
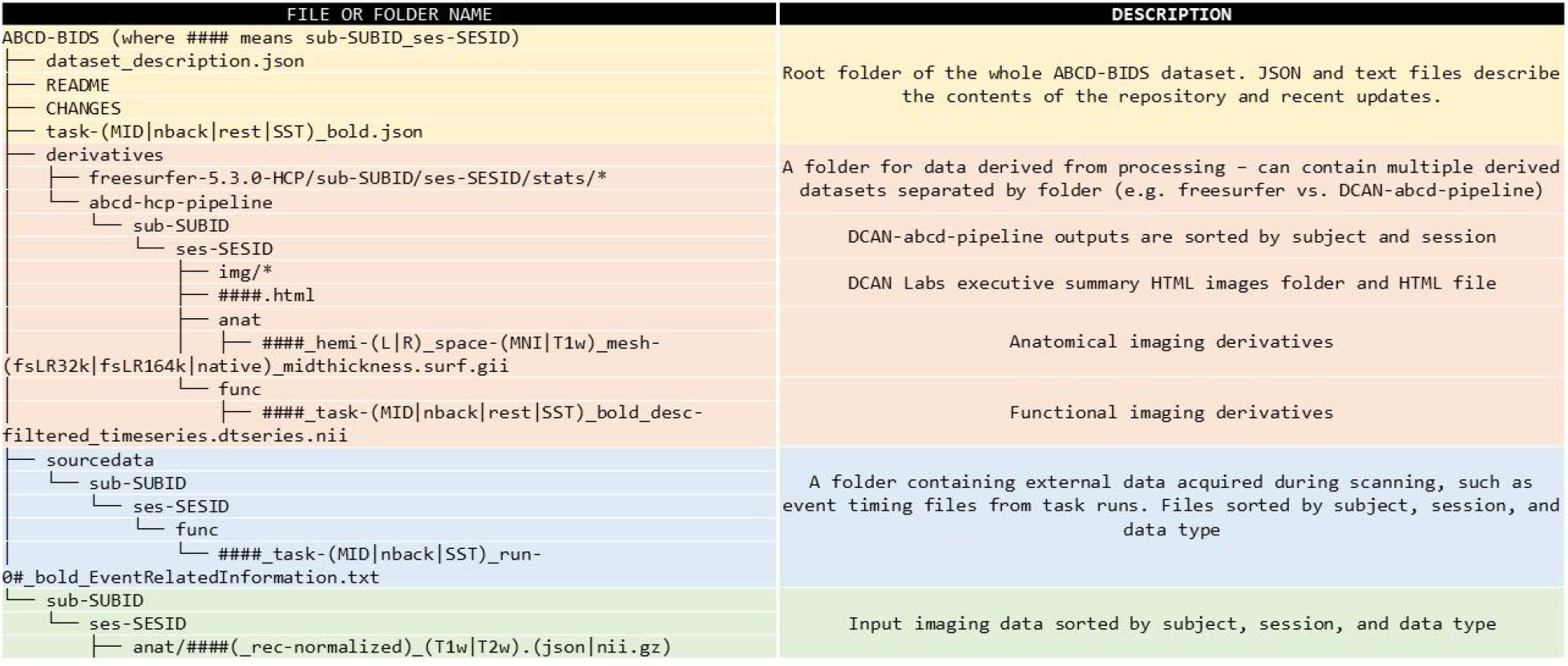
ABCC Share Contents Include a Stable MRI Dataset of 11,877 Children. Subfolder organization in the ABCC (Collection 3165) from top to bottom. **(Yellow)** The 3165 dataset comprises a root folder describing the whole dataset. **(Orange)** Derived data can be found in the derivatives subfolder, organized by pipeline. For example, researchers that want access to derived outputs for further analyses can find them in the “abcd-hcp-pipeline” folder. **(Blue)** Researchers can find additional data, such as the event files needed for task fMRI, in the “sourcedata” subfolder. **(Green)** The BIDS converted input data can be found in each “sub-SUBID” directory for researchers interested in examining data processing strategies.

##### Additional Longitudinal ABCD Data Waves: Stable and Analysis-Ready

The ABCD study is a 10-year longitudinal study with ongoing neuroimaging data acquisition occurring in 2-year intervals. As longitudinal MRI data are acquired in waves, they will be processed with the ABCD-BIDS pipelines and added to the ABCC. As data waves complete processing they will be uploaded and version-stamped with versioned release notes. Therefore, newly uploaded longitudinal data will be stable and analysis-ready. Importantly, data provenance and versioning is currently being implemented with datalad (Hanke et al., 2021).

##### How to Contribute to the ABCC: Community Sharing

In addition to providing data, the ABCC helps facilitate “Open Science” (i.e., open-source and open-data) by enabling researchers to share new, improved, and corrected data derivates along with analytic tools with the broader neuroimaging community. Since neuroimaging research has historically been dominated by small sample sizes (Szucs and Ioannidis, 2020) and BWAS have small effect sizes (Marek et al., 2020; Smith and Nichols, 2018), open resources that allow for data sharing enable independent researchers to leverage bigger data sets to measure effect sizes (Gorgolewski et al., 2017a). However, reproducibility is limited not just by sample sizes, but also by how the measures are derived from a given dataset. For example, processing pipelines can have thousands of different combinations of steps, which can dramatically vary results (Botvinik-Nezer et al., 2020). The parcellation scheme for data analysis may also dramatically impact findings (Wig et al., 2011). The ABCC provides a critically important resource for neuroimaging researchers by enabling them to both share and compare new and old derivatives, such as different processing approaches or parcellations. This shareability enables the scientific community to contrast and compare different approaches, thereby facilitating best practices and higher standards.

Researchers interested in sharing their own derived data can do so through the ABCC. Our NDA-BIDS uploader tool (Earl and Fair, 2021) makes it easy for researchers to prepare and upload their own ABCD data derivatives. Contributed data must undergo an approval process before being uploaded. To request features or additional data to be added to the NDA collection, use this GitHub repository’s Issues page: https://github.com/ABCD-STUDY/nda-abcd-collection-3165/issues. Review and inclusion of data into the ABCC will utilize the community governance structure modified from BIDS as outlined here (https://bids.neuroimaging.io/governance.html). At minimum, the data must be 1) published, 2) BIDS-formatted, 3) the data do not risk reidentification of any participants, and 4) the methods are open source and can be reproduced. After approval, the researcher will be permitted to use the uploader to upload their data to the NDA. This step triggers a formal test and review process, with certifications from both the ABCC and the NDA. Following this process, the data will be available for anyone with ABCC access. Approved data contributors also interested in sharing their code as an ABCC utility can do so by linking their repository to the ABCC open science framework (Feczko et al., 2020a).

### Section II: ABCC Utilities for Reproducible Research

The ABCC utilities enable researchers to evaluate the reproducibility of their findings while minimizing the need to repeat efforts. Here we highlight two utilities included with the ABCC that can be used for assessing reproducibility. 1) The ABCD Reproducible Matched Samples (Cordova et al., 2020a) (ARMS) enables researchers to test the replicability of their findings. 2) Sandwich Estimator for Neuroimaging Data (Feczko et al., 2020b) (SEND) enables researchers to construct robust brain-wide association (BWAS) models that can control for complex repeated measures in big datasets like the ABCD study, while minimizing computing time.

#### ARMS: ABCD Reproducible Matched Samples

##### The Replicability Problem

One of the primary mechanisms by which the scientific community validates a given finding is by repeating that experiment with identical or similar methods in an independent but comparable dataset. The scientific community expects the same or similar results from both datasets. That is, the findings are replicable or reproducible (Goodman et al., 2016). However, it is well-documented in the neuroimaging literature that task and/or rest fMRI results often fail to replicate (for examples see (Botvinik-Nezer et al., 2020; Domínguez-Baleón et al., 2018; Marek et al., 2020; Müller et al., 2017; Uddin et al., 2017).

There are likely several reasons why replicability has been an issue for many studies to date, includinglow signal-to-noise ratio and small effect sizes (Button et al., 2013; Smith and Nichols, 2018)(Button et al., 2013; Marek et al., 2020; Smith and Nichols, 2018)(Button et al., 2013; Smith and Nichols, 2018). Because most studies operate as small sample siloes, publication bias overestimates effects and limits replicability (Marek et al., 2020; Sabuncu et al., 2015; Smith and Nichols, 2018; Walum et al., 2016). The publication system incentivizes findings that meet a “significance” criteria, which is agreed upon by the journal and/or community. As a result, published studies tend to over-represent “significant” findings at small sample sizes but do not publish findings at the same sample sizes that fail to meet this threshold. Therefore, studies based on small sample sizes overestimate the “true” effects due to publication bias. The ABCD dataset, because of its sheer size, provides a ‘best case scenario’ for assuring replicability for many questions relevant to neuroimaging in general and child development in specific.

##### ABCD Reproducible Matched Samples (ARMS) Verify Replicability

Because ABCD comprises a population sample of over 11,877 participants, it can be split into independent “discovery” and “replication” datasets, or in the case of machine learning or model building, train and test sets, where models can be tuned via cross-validation on training datasets and tested on the independent test set. Such an exercise can provide investigators with a training environment to optimize statistical and predictive models and a testing environment to assure findings replicate. Furthermore, by creating stratified groups matched for many factors, researchers can be confident that demonstrations of low replicability likely reflect issues with the measures used and not the samples. Thus, we set out to provide the neuroimaging community with ABCD Reproducible Matched Samples (ARMS). To generate ARMS, ABCD data were split into 3 demographically matched groups (Figure S1): ARMS-1 (N=5,786) and ARMS-2 (N= 5,786) for use as independent datasets, and ARMS-3 (N=305) for template building and model testing. A series of chi-squared tests were used to evaluate matching for each demographic variable.

To create the matched ARMS, we chose 9 salient sociodemographic factors thought to be important for developmental outcomes (site, age, sex, ethnicity, grade, highest level of parental education, handedness, combined family income, and exposure to anesthesia), and accounted for family structure (Marek et al., 2019). Anesthesia exposure was included as a matching variable to account for differences in major medical interventions and the possible effects on behavioral and neurodevelopmental outcomes (Schneuer et al., 2018). To maximize the relative independence of the two datasets, family members were kept together in the same ARM and the groups were matched to have equivalent numbers of sibling and twin pairs, and triplets.

Comparisons across site, age, sex, ethnicity, grade, highest level of parental education, handedness, combined family income, exposure to anesthesia, and family-relatedness show no significant differences between ARMS-1 and ARMS-2 (Table 2; Table S2). Comparisons of the counts and means for each of these factors show that ARMS-1 and ARMS-2 are almost identical samples. Gender shows the largest absolute difference of 2.5 percent, and no other demographic variables differ by more than 1 percent (See supplemental materials: Figures S3; S4 for more details). Although not explicitly matched, cognitive performance was identical across ARMS (see supplemental materials: “Experiment II methods”).

**Table 2.**
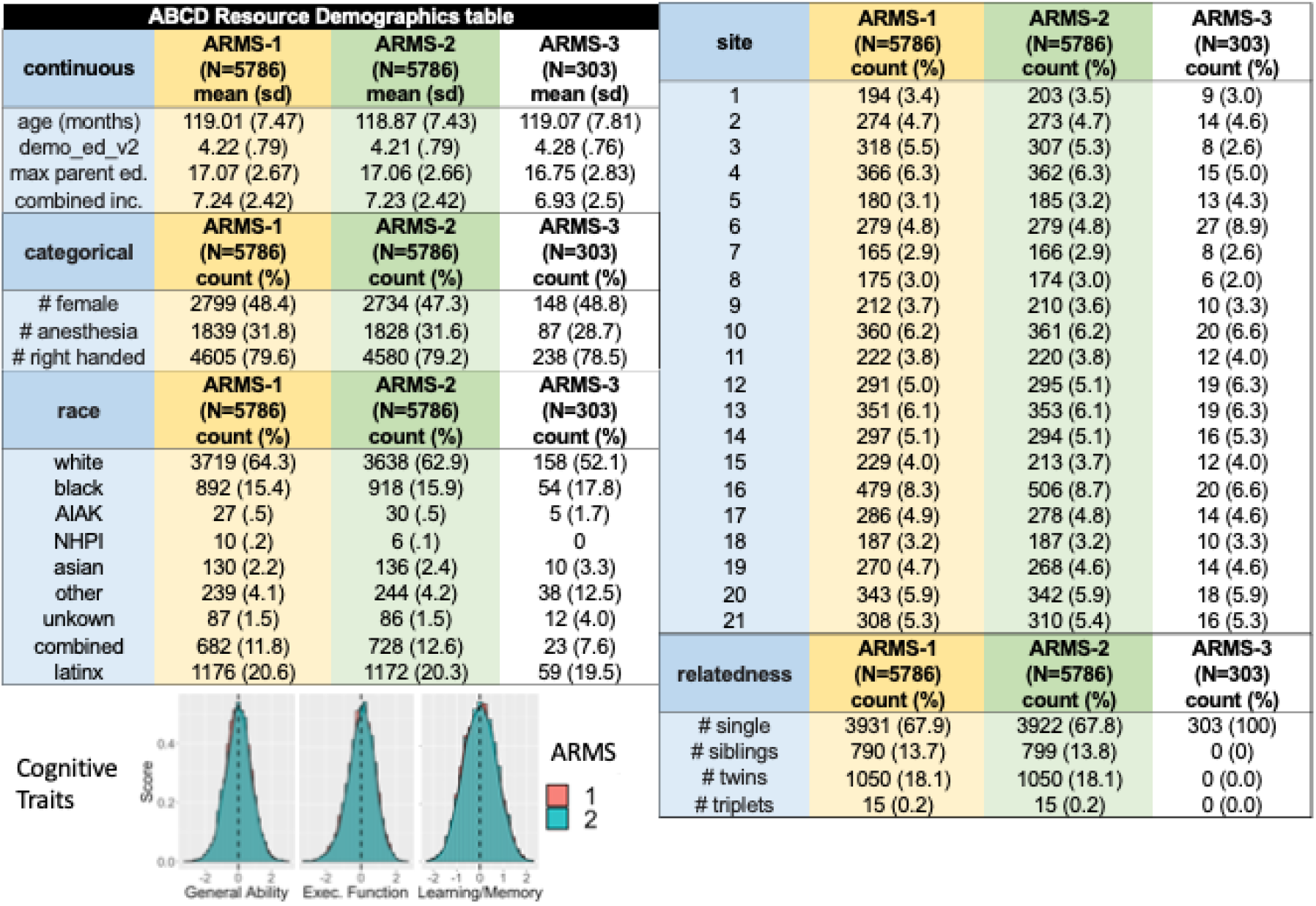
ARMS-1 and ARMS-2 Matched for Demographic Variables. Demographics for the MRI datasets are depicted in this table. Differences between ARMS-1 and ARMS-2 are minimal, with proportion white (race) showing the largest difference at 1.4 percent. No other factor shows differences greater than 1 percent. ARMS-3 differences are larger for both site and race but not other demographic factors, additionally, ARMS-3 does not include related family members. A graph of cognitive traits shows no differences between ARMS-1 and ARMS-2 for General Ability, Executive Function, or Learning and Memory.

While ARMS provides an exciting approach to measuring replicability for data analyses, such as BWAS (see section III below), we do note that inferences regarding replicability may be limited compared to a true independent replication, or generalization to a different study. In addition, some findings may not replicate with ARMS simply because the effect sizes are so low that a dataset larger than the split half are required to identify the true effects. If this is the case, however, investigators should proceed with caution as even the full sample may not be large enough for characterizing the true effect of a given analysis (genetics citation maybe).

Altogether, the ABCD dataset in combination with ARMS provides investigators with the tools to generate reliability and reproducibility estimates for findings obtained from the ABCD Study, which are further explored in section III.

#### Sandwich Estimator for Neuroimaging Data (SEND): Robust BWAS

##### Complexities of Nested Covariates in Multi-site Studies

Brain-wide association studies (BWAS) often must take into account covariates that may impact an analysis of interest. For example, a study of brain volume may incorporate a measure of intracranial volume as a covariate due to an independent association between head size and brain volume. Although the ABCD study contains a diverse sample, the way in which it was acquired causes normally independent covariates to become “nested”, or structured. For example, the ABCD sample was collected across 21 research sites (Casey et al., 2018; Volkow et al., 2018), and MRI data were acquired across three different platforms (GE, Siemens, Philips). As a result, both site and platform effects are likely to impact the measured data. Because not all dependent measures are overlapping (e.g., not every site has a different scanner platform), the covariance structure of the data is considered to be nested.

In smaller data sets, nested covariance structure could be accounted for with linear mixed effects models. However, in neuroimaging datasets where each image itself comprises tens of thousands of data elements, the sheer size and complexity of the ABCD study requires computationally intensive analyses – in some cases, statistics over a billion data points. On such datasets using a single computer or compute node, linear mixed effects models can take more than six months to complete (Guillaume et al., 2014). In addition, properly adjusting statistical significance in this scenario (e.g., when using permutation testing (Eklund et al., 2016)) becomes less exact as the number of tested variables increases. Therefore, we developed a marginal model approach (Guillaume et al., 2014) and a software package for testing brain behavior associations in large-scale studies such as ABCD - the Sandwich Estimator For Neuroimaging Data (Feczko et al., 2020b) (*SEND)*. A marginal model approach can be nearly as efficient as a mixed model without requiring slow iterative optimisation; the only potential shortcoming of a marginal model is that you cannot estimate and make inference on random effect variance components. However, these effects are implicitly accounted for and, as we generally are not interested in these terms, it is not a significant shortcoming.

##### SEND: Accounting for Nested Variables Efficiently

SEND combines an ordinary least squares (OLS) model with a sandwich estimator (SwE). An ordinary least squares (OLS) model is a simple linear model with population-level covariates (e.g. “age”, but not subject-specific slopes for age) parameters, where the parameters are estimated via least squares. A sandwich estimator (SwE: see supplemental materials for details -- SI Figure 5), measures the nested covariance structure and estimates the standard error accounting for this structure(Vandekar et al., 2019). Even though the OLS model is ignorant of any nested structure, the regression coefficients are unbiased, and for sufficiently large samples so is the sandwich estimator of variance (Eicker, 1963). Combined, these steps can be solved algebraically (see: Supplemental Materials: Marginal Framework), so they are faster than traditional linear mixed models (Figure 2C).

Statistical significance is tested via a wild bootstrap instead of a permutation test. Permutation testing (Eklund et al., 2016; Winkler et al., 2016) requires exchangeability under the null hypothesis, something in general violated by the nested data (Anderson and Braak, 2003). A wild bootstrap (see: Supplemental Materials: Marginal Framework) enables users to properly control for false positives. As an alternative to a permutation test, the wild bootstrap multiplies each individual’s residual by randomly resampled numbers reflecting the structure of the data; for example, every residual within a nested grouping receives the same multiplier. For each bootstrap, the new null data are composed as the sum of the original fitted data and the resampled residuals, and the marginal model is built using the new data. Such a procedure enables one to construct a series of null models(Vandekar et al., 2019) and assess the statistical significance of MRI findings using traditional cluster (Winkler et al., 2016), threshold-free cluster (Smith and Nichols, 2009), or enrichment (Eggebrecht et al., 2017) analyses.

Figure 3 depicts the SEND package and its outputs. The analysis comprises two parallel workflows. ConstructMarginalModel (Figure 3A) performs the analysis on the observed data. A linear model is fit for brain imaging and external measures (Figure 3A; top), statistical maps then undergo statistical testing, such as via traditional cluster (Winkler et al., 2016), threshold-free cluster (Smith and Nichols, 2009), or enrichment (Eggebrecht et al., 2017) analyses (Figure 3A; middle). Brain imaging inputs can be encoded as NIFTIs, GIFTIS, or CIFTIs (Figure 3A; bottom). If the MRI inputs are surfaces or volumes, ConstructMarginalModel can perform cluster detection. If the inputs are connectivity matrices, ConstructMarginalModel can perform an enrichment analysis (Cordova et al., 2020b; Eggebrecht et al., 2017; Feczko et al., 2018; Marek et al., 2019) to determine network effects (e.g. effects on the visual system or between the default and somatomotor networks; Figure 3A; bottom) as opposed to statistics on individual connections.

**Figure 3.**
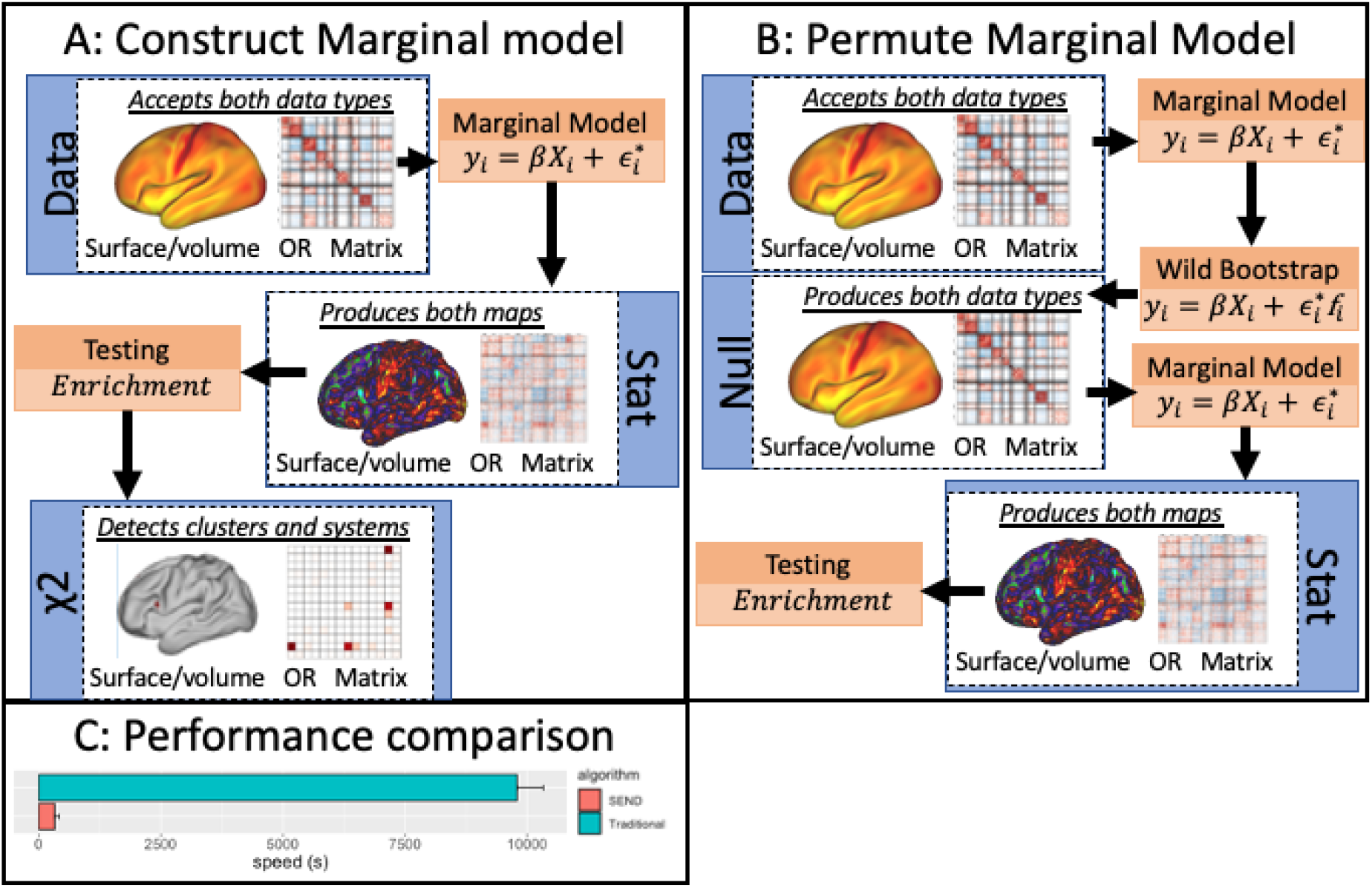
SEND workflow. The SEND package comprises two tools for conducting significance testing on ABCD data. **(A) ConstructMarginalModel** fits a marginal model to input data, which may comprise surface metrics like cortical thickness, volumes, or connectivity matrices. Statistical maps are computed and then z-score normalized before performing testing, which may represent cluster detection or an enrichment analysis. **(B) PermuteMarginalModel** uses a wild bootstrap procedure to simulate a series of datasets under the null hypothesis. Each dataset undergoes the same modelling as **ConstructMarginalModel**, to construct a null distribution of the final test statistic. **(C) SEND** performance speed on 10 iterations using 3000 subjects on ~62,000 tests compared to traditional performance via Geepack. On the same dataset, Geepack takes ~10,000 seconds to compute test statistics, whereas SEND takes ~300 seconds.

To assess significance, PermuteMarginalModel constructs the null distribution using the wild bootstrap (Figure 3B). It first calculates the fitted model on the observed data from ConstructMarginalModel (Figure 3B; top). Using the residuals and fitted values from the model, PermuteMarginalModel multiplies each case’s residual by a number resampled from a random symmetric distribution (Figure 3B; middle). Several different distributions can be selected (e.g. Radenbacher), the default of Radenbacher is based on previous recommendations for best standards and practices to limit false positives(Guillaume et al., 2014). The multiplied residuals are added back to the fitted values to form a bootstrap sample. PermuteMarginalModel then calculates the same model as ConstructMarginalModel, but only saves the final test statistic (Figure 3B; bottom).

We compared performance speed using SEND to Geepack (Halekoh et al., 2006), which allows for the same mixed and marginal modelling procedures described above. We performed 10 iterations of both analyses, using the identical parameters described in Section III, example 2. As shown (Figure 3C), SEND performs its calculations 30 orders of magnitude faster than the traditional R package.

##### SEND is Flexible and Easy-to-use

By devising the workflows in parallel, a full SEND analysis is scalable for any system design with minimal effort, from a single computer to a batch submission to a cluster, to cloud-based services. With cluster or cloud computing, researchers can potentially run thousands of permutations in hours instead of days.

The marginal model package can be installed from within R and is available via GitHub (Feczko et al., 2020b). The software accepts standard NIFTI, GIFTI, or CIFTI inputs, which are well documented neuroimaging formats, and therefore can model outputs from other imaging pipelines. Examples, documentation and parameter files are provided so that novice R users can perform analyses by running SEND on the command line. Advanced documentation is provided for expert users. The section below highlights how the ABCC can be used to examine reproducibility of the ABCC dataset.

### Section III: Sample Size Considerations with Big Data like ABCD

In this section, we use the utilities described in Section II to examine the reproducibility and reliability of ABCC parcellated resting state functional connectivity matrices. Similar analyses of cortical thickness measurements and other morphological measures in the ABCC are also provided in supplementary material (see: “Experiment I”). Before we move forward, it is important to clearly define terms as used in this section. (Note: For the remainder of this manuscript we will use the terms reproducibility and reliability to denote different approaches used, though we recognize that such terms can be defined in multiple ways (Goodman et al., 2016)).

#### Reproducibility

The similarity of BWAS test statistic patterns (i.e. the patterns of association RSFC and cognitive ability associations) between two split-partitions. Specifically we use the ARMS and their subsets to independently calculate a given measure or test statistic and compare the patterns via spatial correlation (See: supplementary materials “Experiment I” for a description of the methods).

#### Reliability

The maximum effect size observed from group comparisons between subset samples to a fully independent but matched subset sample. (See: supplementary materials “Experiment I” for a description of the methods). Others may be more familiar with the term “test-retest reliability”, which is not measured here.

We focus on reproducibility and reliability to highlight: 1) That neuroimaging analytics requires large samples for reproducible and reliable findings, and 2) that even with large samples, brain-behavior association reproducibility will vary depending on the level of noise in the data. In short, we show that the data are reproducible and relatively stable once the sample size exceeds a thousand subjects. Reproducibility and reliability are reduced with smaller samples such that group comparisons may be unstable. We also show that differences in reproducibility will vary as a function noise in the data (for both dependent and independent variables).

### Example 1: Between-Group Comparisons

Our first example examines the within-study reproducibility, and reliability of functional connectivity matrices. The full analyses are provided in the supplementary material for other neuroimaging modalities (e.g. sulcal depth) and data trimmed to 5 minutes instead of 10 minutes (Figures S7-S16). Subset reproducibility and reliability analyses can go in two directions - using ARMS-1 as the baseline or ‘gold standard,’ or using ARMS-2 as the gold standard. In the main manuscript, we show the latter, however, the same finding is replicated for the former in supplementary materials (Figures S10, S13).

#### Reproducibility via ARMS

ARMS was utilized to measure the mean and inter-subject variance reproducibility for connectivity matrices and mean morphological measurements. Reproducibility of group average connectivity matrices was conducted by calculating the average ARMS-1 and ARMS-2 matrices and computing the correlation and difference between ABCD -1 and ARMS-2. Reproducibility of group variance was conducted in a similar fashion. We also measured the correlation between mean cortical thickness (Figure 4), folding curvature, sulcal depth, and myelin (see supplemental materials for a detailed discussion; Figures S11-S16).

**Figure 4.**
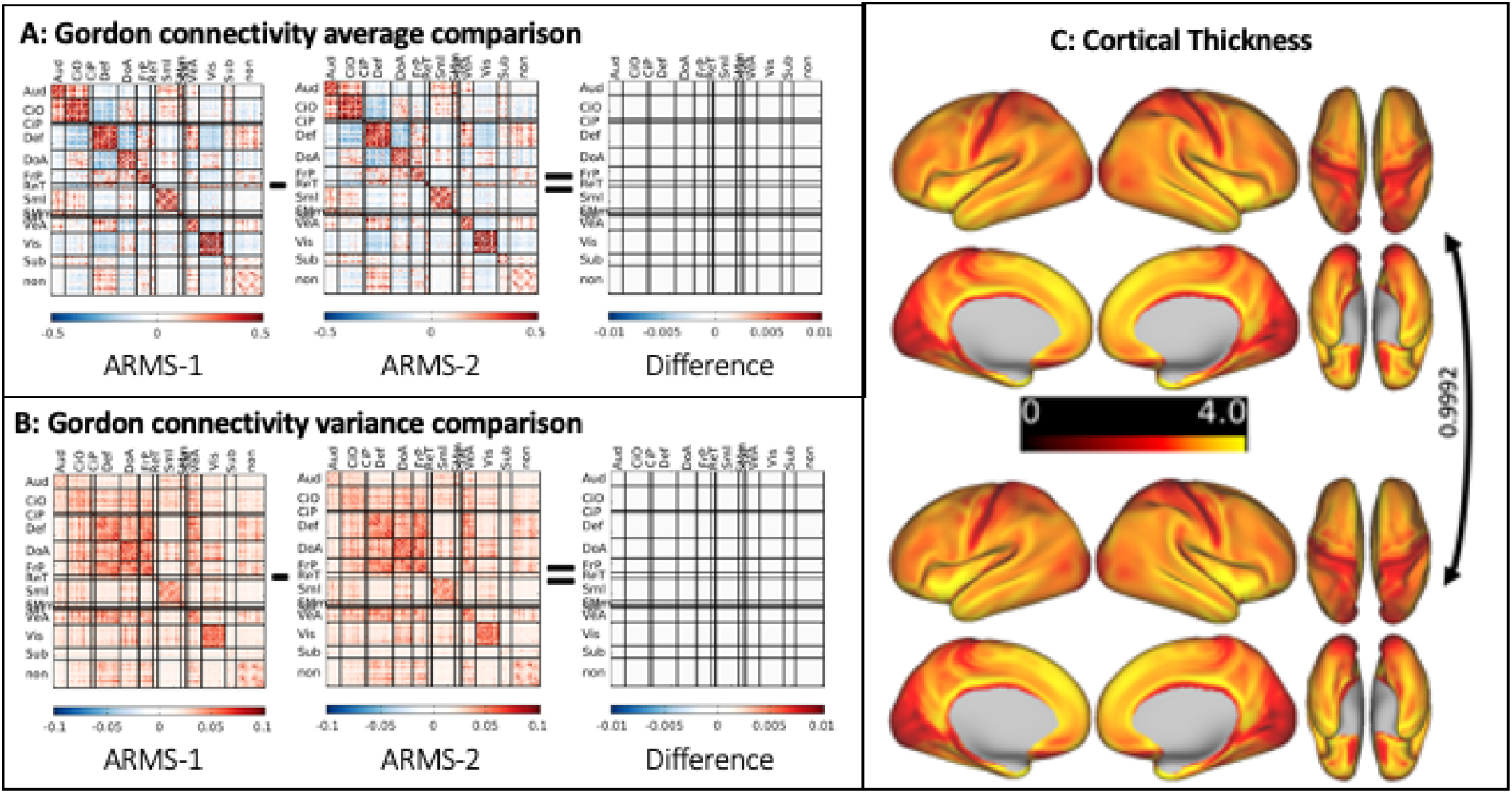
Functional Connectivity Matrices Are Reproducible between ARMS Datasets (ARMS-1, ARMS-2). **(A)** Mean connectivity matrix for ARMS-1 (left) and ARMS-2 (middle) and the difference between them (right). Correlations are sorted by the Gordon-identified community structure. The proportion of variance explained is indicated by the arrow in between the two group matrices. **(B)** Variance connectivity matrix for ARMS-1 (left) and ARMS-2 (middle) and the difference between them (right). Correlations are sorted by the Gordon-identified community structure. The proportion of variance explained is indicated by the arrow in between the two group matrices. **(C)** Cortical thickness compared between ARMS-1 (top) and ARMS-2 (bottom). Spatial correlation between ARMS-1 and ARMS-2 shows near identical reproducibility (r^2^=0.99).

When matching ARMS-1 and ARMS-2 for the highest data quality and amount of time points available, precisely 10 minutes of data were selected per subject (i.e. “trimming”: see supplemental materials for details). Mean matrices from ARMS-1 and ARMS-2 show high within-study reproducibility. Point estimates for ARMS-1 (Figure 4A; left) and ARMS-2 (Figure 4A; middle) were nearly identical for 10 minutes of data (r^2^=0.9999; Figure 4A; right). Similarly variance estimates for ARMS-1 (Figure 4B; left) and ARMS-2 (Figure 4B; middle) were nearly identical for 10 minutes of data (r^2^=0.9922; Figure 4B; right).

#### Subset reproducibility

Subset reproducibility was measured at sample sizes of 2, 3, 4, 50, 75, and 100. As with full-group reproducibility, we calculated the average connectivity matrix for the ARMS-1 subset and computed the correlation to the average ARMS-2 full matrix at sample sizes of 2, 3, 4, 50, 75, and 100. We chose such sample sizes because the mean matrix reproducibility plateaus quickly with a small set of subjects. We also compared the variance matrix reproducibility as well, but calculated the correlation between them at larger sample sizes (26, 40, 50, 75, 100, 200, 300, 400, 500, 750, 1000, and 1250), because much larger samples are needed for reproducible variance matrices.

Subset reproducibility began to reach correlations of 0.99 for the mean matrices with as little as 20 subjects (Figure 5A), but sample sizes over 1000 were needed to achieve the same level of reproducibility (Figure 5B). Practically speaking, this is a consequence of the variance estimates (Figure 4B) being smaller than the mean estimates (Figure 4A). Simulations of random sets of variables for two random samples show the same pattern (Figure 5D), which flips when the variance is greater than the mean values (Figure 5E). At smaller sample sizes, greater inter-subject variability in the central tendency and distribution of data means that the sample is not representative of the underlying population, even with demographic matching. Such sampling variability may drive misestimation of effect sizes when performing phenotype/BWAS studies, as seen below.

**Figure 5.**
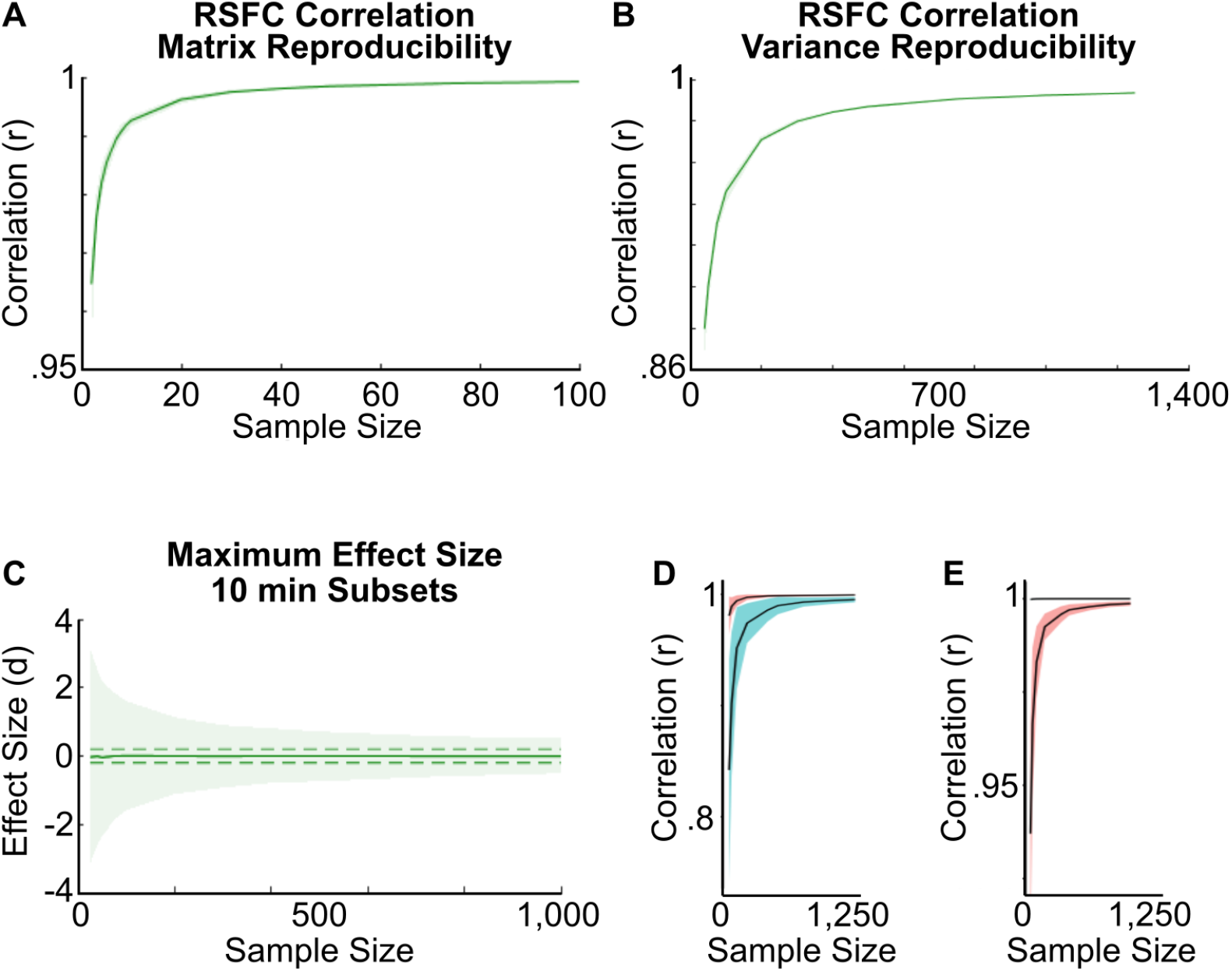
Subset Reproducibility for Variance and subset reliability for Effect Sizes Requires Over 1000 Subjects. **(A)** Subset reproducibility analyses for ARMS-2 subsets to ARMS-1 for mean connectivity matrices. X-axis represents the sample size while Y shows the correlation. **(B)** Subset reproducibility analyses for ARMS-2 subsets to ARMS-1 for variance connectivity matrices. X-axis represents the sample size while Y shows the correlation. **(C)** The subset reliability analysis measuring maximum cohen’s d effect sizes for group comparisons between ARMS-2 subsets to ARMS-1 for 10 minutes of data. Here, we calculated a simple cohen’s d, calculated as the difference of means divided by the pooled standard deviation. X-axis depicts the sample size while Y shows the effect size per connection per subset. **(D)** Simulations of random sets of variables with means (black line with 95 and 5 percentiles shaded in red) of 0.3 +/− 0.1 and standard deviations (black line with 95 and 5 percentiles shaded in aqua) of 0.1 +/− 0.2. The x-axis depicts the sample size while the y-axis depicts the correlation. **(E)** Simulations of random sets of variables with means of 0.3 +/− 0.6 and standard deviations of 0.1 +/− 0.03. Graph is the same as **(D)**.

#### Effect size reliability

Finally, we conducted a comparison between ARMS-1 and ARMS-2 and calculated an effect size measuring group differences across all connections per subset. Here, we calculated a simple cohen’s d, calculated as the difference of means divided by the pooled standard deviation. This analysis enables us to estimate the possible effect sizes observed under the null hypothesis for group comparisons. The results also help guide researchers towards potential effect sizes for group comparisons as a function of sample size. Because the subset reliability analyses produce stratified matched subsets, we anticipate that the “true” effect size of group subset comparisons are zero; the differences between the mean matrices are zero (Figure 4A). The subset reliability analysis shows that observed effect sizes move closer to zero as sample size increases (Figure 5C); however even with 1000 participants, for between-group comparisons of single connections, the “true” effect size may be misestimated by as much as a cohen’s d of 1 (Figure 5C).

Taken together, these analyses strongly suggest that effect size estimates of single connections may require thousands of subjects when performing whole-brain analyses. Additional analyses and similar findings using other modalities supporting these conclusions are shown in supplemental materials, and highlight the importance of carefully considering realistic power estimates and designing analyses with ABCD data with an eye toward large samples for group comparisons. Because even cohen’s d of 0.5 can occur with samples with hundreds of subjects, sample sizes < 1000 for many univariate group comparisons using ABCD imaging data are likely to be insufficient.

### Example 2: Brain-Wide Association Studies (BWAS)

Our second example conducts three different analyses of brain-behavior associations to illustrate the intersection of parameter and measurement selection. As an example, we used the three cognitive traits extracted from the ABCD cognitive task data ((Luciana et al., 2018); see supplemental materials for details) validated in a prior report (Thompson et al., 2019). Specifically, we performed BWAS using the three cognitive traits identified from the ABCD cohort: general ability (PC1), executive function (PC2), and learning/memory (PC3). To avoid potential bias and inflation of effects (Dinga et al., 2019; Sripada et al., 2019), it is important to treat ARMS-1 and ARMS-2 samples as purely independent. Therefore, the traits were extracted from ARMS-1 and ARMS-2 independently. Results showed strong reproducibility of each trait across the two datasets (Table 2) and the original report by Thompson *et. al.* (Figure S6). PC1 is shown in the main manuscript (Figure 6), while additional analyses for PC1 (enrichment), PC2 and PC3 figures can be found in the supplemental materials (Figures S17-S22). For simplicity, we will discuss the subset reproducibility analyses from ARMS-2 subsets (5- or 10-minute) to the full ARMS-1 (baseline or gold standard) 10-minute dataset. We replicated the same findings from ARMS-1 subsets to the full ARMS-2 10-minute dataset, which are discussed in supplemental materials (Figure S22).

**Figure 6.**
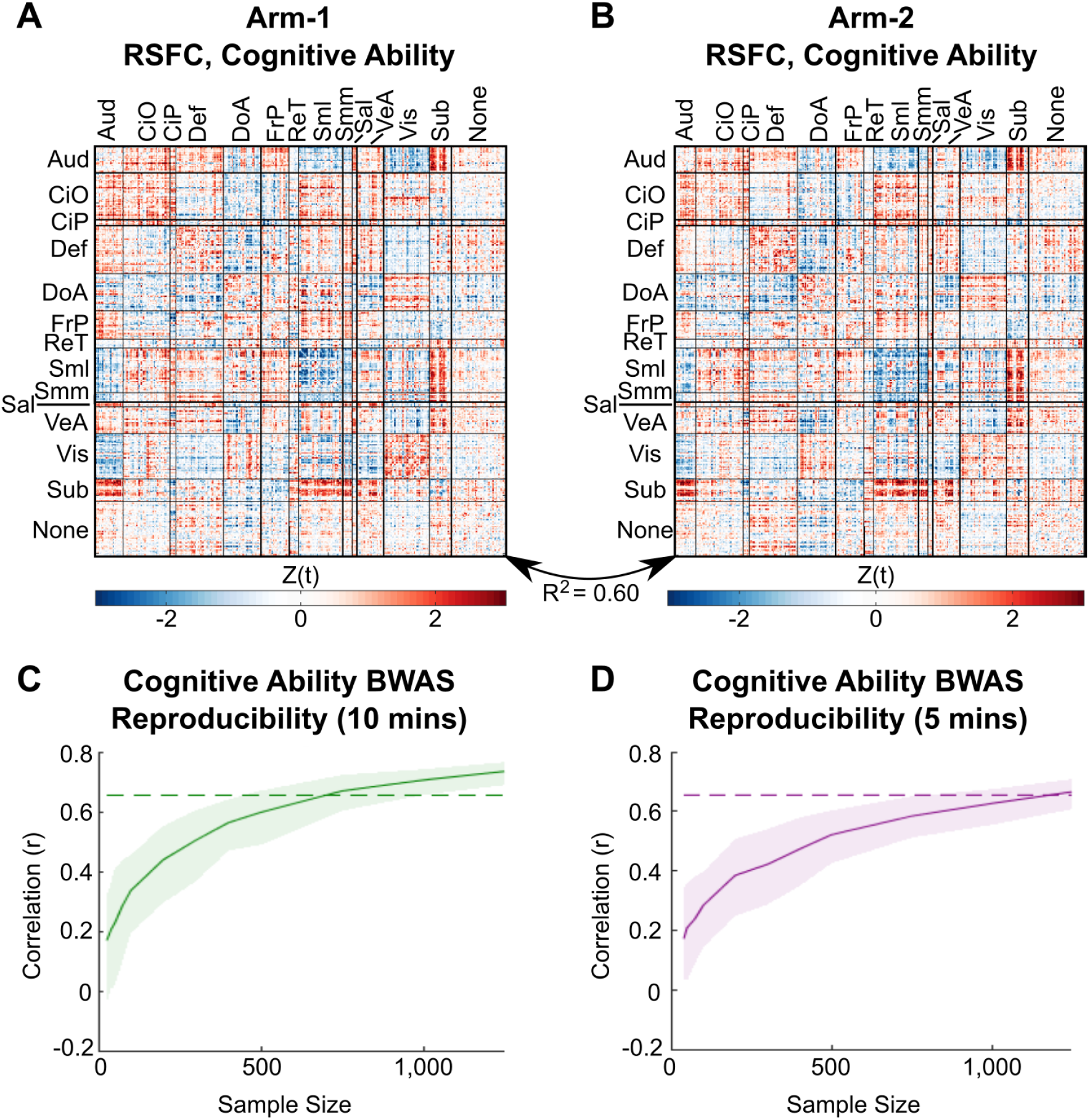
Brain Behavior Associations between Functional Connectivity and General Cognitive Ability. **(A)** Matrix of Z-scores for parcellated connections for ARMS-1. Rows and columns represent ROIs sorted by network assignments, each cell contains the z-score test statistic for general ability predicting the given connection (FC ~ PC1). **(B)** Z-score matrix for ARMS-2. The proportion of variance explained between the two maps is indicated by the double-arrow curve. **(C)** Plot of the correlation between subsets from ARMS-2 to the full ARMS-1 using 10 minutes of data, showing the reproducibility of RSFC patterns associated with cognitive ability. X-axis depicts sample size while Y shows the correlation between subsets. The dotted line indicates the threshold where correlations exceed 80 percent of the correlation from the full comparison. **(D)** The same plot shown for subsets using 5 minutes of data instead of 10.

#### Reproducibility of RSFC patterns measured by ARMS

Brain-behavior associations for each trait were computed on the 10-minute functional connectivity matrices (Gordon parcellation) used in ARMS via the marginal model and an enrichment analysis (SEND). For all three cognitive traits, per connection, a marginal model was fit with the given trait as a predictor and the connectivity as the outcome (FC ~ PC). Site and gender were included as nested variables to control for batch effects. After fitting a marginal model at each connection, standardized estimates were normalized by converting them to z-scores.

For each trait, we performed the above analyses for both ARMS-1 and ARMS-2 separately (i.e. via ARMS). We examined the within-study reproducibility of the mass-univariate tests by measuring the correlation between ARMS-1 and ARMS-2 connectivity matrix z-score maps. Because differences between statistical maps may not be sufficiently captured by correlations, we also visualized the differences between ARMS-1 and ARMS-2 by subtracting one map from another.

The correspondence of ROI-based connectivity with general ability (PC1) in ARMS-1 (Figure 6A) and ARMS-2 (Figure 6B), showed robust within-study reproducibility (PC1; r^2^=0.67). Differences between the statistical maps were small on average (0 +/− 0.57). Notably, connections within the somatomotor lateral (SMl), and between the subcortical (sub) and SMl/auditory(aud) systems show higher enrichment of z-scores associated with general ability (see supplemental materials). These findings are consistent with recent large-scale studies within the ABCD data set (Marek et al., 2019), and from other large transdiagnostic samples using different processing strategies and different parcellation schema (Chen et al., 2020). Importantly, however, while reliability and reproducibility were relatively strong for general ability (PC1; r^2^=0.67; Figure 5), it was less so for executive function (PC2; r^2^=0.37; SI figure S18) and learning/memory (PC3; r^2^=0.46; SI Figure S20). This dynamic simply highlights that with weaker effect sizes, even more subjects are required to reach stability.

#### Subset reliability

Recall that reliability is defined as the similarity of test-statistics between a subset of one sample to an independent, matched, full sample. Therefore, we examined the effect of sample size on brain-behavior association reliability (see: supplemental materials for more details). The analysis was conducted on sample sizes of 26, 40, 50, 75, 100, 200, 300, 400, 500, 750, 1000, and 1250. Per sample size, 100 subsets were generated from ARMS-1 and ARMS-2 separately. Per subset, FC-trait z-scores were derived via marginal models. The correlation between each subset’s FC-trait z-score matrix was conducted against the other group’s FC-trait z-score matrix produced on the full 10-minute dataset.

Notably the within-study reliability increases as a function of sample size (Figure 3D). Perhaps somewhat concerning is that to reach 80 percent of the maximum reproducibility observed (r^2^=0.54) more than 1000 participants are required (Figure 3D: grey dotted line) – a sample size rarely achieved for most studies.

Selecting only 5 minutes of data to generate individual matrices reduced reliability (Figure 3E). Overall, the sample size needed to achieve similar RSFC patterns reproducibility to the 10-minute dataset was reduced (Figure 3E). Even with 1,250 subjects, the correlation between subsets does not achieve 80 percent of the maximum (Figures 3E). In fact, it takes 1,250 subjects with 5 minutes of data to achieve the same reproducibility as 750 subjects with 10 minutes of data (Figures 3D), highlighting the importance of maximizing the percentage of low-movement data collected (Dosenbach et al., 2017).

#### Additional Considerations

Despite the reproducibility and reliability of the findings above, one needs to be cautious regarding causal interpretations regarding mechanism (Mehler and Kording, 2018). Any BWAS, no matter how significant, suffers from the heterogeneity problem (Feczko and Fair, 2020; Feczko et al., 2019; Satterthwaite et al., 2020), and therefore may be mediated both by important factors, such as genetic expression or inflammatory markers, and unimportant factors, such as head motion. Furthermore, it is possible that the reproducible effects here are additive, and therefore better represented by a multivariate model, such as the bayesian polyvertex estimator(Zhao et al., 2021). While the work above shows a reproducible association between general ability and functional connectivity, more work is needed to determine *why* this association is reproducible (Pearl and Mackenzie, 2018).

## Conclusion

### ABCC-Data and Utilities Promote Accessibility and Security

The ABCC-data (ABCD-3165) (Feczko et al., 2020a) and utilities enable researchers to study child brain development and mental health with maximum accessibility and security. The difficulties of big data management make combinations of data and utilities critical for researchers to perform fast and robust analyses easily. Such combinations should be harmonized, so researchers can focus less on data management, curation, and processing and more on best practices and standards for their analyses.

#### Child Mental Health Research Needs Secure and Accessible Consortia

As neuroimaging research increases in scale, due to the need for large samples, large amounts of PHI and imaging data will be acquired across multiple sites in consortia like the ABCD study. While such a scale is necessary for reproducible and beneficial research, it is not sufficient. Governance of the data via institutions like the NIMH Data Archive (NDA) is needed to limit both privacy risks, such as re-identification, and researcher accessibility; researchers cannot upload individually-derived measures to the current collection (ABCD-2573). The ABCC (ABCD-3165) provides this governance structure but also enables researchers to share new individualized measures (or derivatives), such as new brain parcellations. The combination of both types of collections maximizes security and accessibility for ABCD data.

#### With Large Consortia, Researchers need Fast, Easy, and Robust Utilities

The ABCC-data (ABCD-3165) alone is not sufficient for researchers to quickly incorporate ABCD study data into their own analyses. The sheer size of the data requires researchers to plan storage, computational, and software development/implementation costs simply to use the data itself. Applying these ready-to-use ABCC utilities alongside the ABCD-data streamlines the reformatting and processing steps, allowing researchers to spend more time planning analyses and actually using the data. Each utility serves a different purpose within a given neuroimaging analysis and can be run independent from the ABCC. The ABCD-BIDS pipeline enables researchers to reproduce ABCD-BIDS style processing. Stratification utilities like ARMS enable researchers to directly test whether their findings replicate in independent matched samples, providing confidence for more robust findings. Analytic tools like SEND enable researchers to quickly and robustly construct linear models for neuroimaging data that also control for the “nested” structure of ABCD data. While the ABCC-data and utilities are highly flexible, running them together is straightforward.

#### Research Benefits Most When Data and Utilities Work in Harmony

The ABCC makes the ABCD data accessible to more researchers and simplifies collaborations. In the examples provided above, we show how the ABCC-data and utilities can be combined to address considerations regarding sample size and data processing. For the ABCC, functional connectivity using a standard parcellation is highly reproducible, but only stabilizes with large sample sizes of at least a thousand subjects. Furthermore, brain-wide associations can vary in their reproducibility and reliability depending on the trait selected, even if both traits are measured from the same set of behaviors. Therefore, 21st century mental health research depends on leveraging big data, maximizing both security and accessibility, and the ABCC can help facilitate both of these goals for the pediatric research community.

## Acknowledgments

We would like to thank Anders Dale for his thoughtful comments on this manuscript. We thank the families who have participated in this research. We are grateful to Susan Tapert, Ph.D. who has expertly guided the work of ABCD’s assessment workgroups, as well as Margie Mejia-Hernandez for her support to the ABCD Workgroup on Neurocognition. Data used in the preparation of this article were obtained from the Adolescent Brain Cognitive Development (ABCD) Study (https://abcdstudy.org), held in the NIMH Data Archive (NDA). This is a multisite, longitudinal study designed to recruit more than 10,000 children age 9-10 and follow them over 10 years into early adulthood. The ABCD Study is supported by the National Institutes of Health and additional federal partners under award numbers U01DA041048, U01DA050989, U01DA051016, U01DA041022, U01DA051018, U01DA051037, U01DA050987, U01DA041174 (B.J.C., R.M.), U01DA041106, U01DA041117, U01DA041028, U01DA041134, U01DA050988, U01DA051039, U01DA041156, U01DA041025, U01DA041120, U01DA051038, U01DA041148 (B.N., H.G., R.W., D.A.F.), U01DA041093, U01DA041089, U24DA041123 (B.J.C.), U24DA041147 (H.G.). Dr. Elizabeth Hoffman was substantially involved in all of the cited grants consistent with her role as a Scientific Officer. A full list of supporters is available at https://abcdstudy.org/federal-partners.html. A listing of participating sites and a complete listing of the study investigators can be found at https://abcdstudy.org/consortium_members/. This manuscript reflects the views of the authors and may not reflect the opinions or views of the NIH, its affiliated Institutes, Centers or offices, or ABCD consortium investigators. The ABCD data repository grows and changes over time.

This work was also supported by NIH grants MH100019 (S.M.), MH121518 (S.M.), DA007261 (D.F.M.), MH112473 (T.O.L.), MH115357 (D.A.F.) and a diversity supplement (MH115357-02S1 (R.H.), MH096773 (D.A.F, N.U.F.D.), MH122066 (D.A.F., N.U.F.D.), MH121276 (D.A.F., N.U.F.D), MH124567 (D.A.F., N.U.F.D.), R01MH113550 (T.D.S.), R01MH112847 (T.D.S), RF1MH116920 (T.D.S.), R01MH120482 (T.D.S), NS088590 (N.U.F.D.), and the Andrew Mellon Predoctoral Fellowship (B.T.-C.), Lynne and Andrew Redleaf Foundation (D.A.F.), Kiwanis Neuroscience Research Foundation (N.U.F.D.), the Jacobs Foundation grant 2016121703 (N.U.F.D.).

