## Supplemental Materials for "Adolescent Brain Cognitive Development (ABCD) Community MRI Collection and Utilities"

### *SI Material*

The supplemental materials comprise three additional sections that enrich the points discussed in the main manuscript.

#### Section II

##### Additional modifications for ABCD BIDS pipeline

###### 1) PreFreesurfer changes

The HCP PreFreeSurfer stage takes a set of anatomical T1-weighted and T2-weighted raw images, with an optional field map, and computes several features: a linear transform to the standard template (each), a brain-ROI mask, relevant image restoration, and a nonlinear registration to the standard atlas using FSL FNIRT. As this is the first stage, any small errors in extracting these features can cause unexpected problems in downstream preprocessing stages. We modified the pipeline to permit an optional T2w image, allowing legacy datasets to be processed. The added benefit of an optional T2 may allow for greater data inclusion when a high quality T1 but poor quality T2 were acquired, which may affect the MRI processing and resultant imaging data quality. We also perform the nonlinear registration to the standard atlas in 3) PostFreeSurfer, which increases the effectiveness of the registration.

###### 2) Freesurfer changes

The HCP FreeSurfer stage is kept identical in function, performing semantic segmentation of brain structures, reconstruction of the white/gray matter surface boundary and the gray matter/pial surface boundary, and surface registration using surface feature maps. The additional HCP process of improving gray matter/pial surface estimates by using the T2 weighted image is made optional based on the presence of a T2w image.

###### 3) PostFreesurfer changes

The PostFreesurfer stage converts freesurfer outputs into the standard CIFTI space. Modifications to this stage better harmonize the reconstructed surfaces and corresponding metrics. As discussed in 1) PreFreesurfer, nonlinear registration to the standard MNI atlas is computed using the refined brain mask acquired through 2) Freesurfer. This brain mask, now refined by the reconstructed surfaces, provides better alignment to the standard template than the preliminary mask used in 1) PreFreesurfer. Additionally, the ANTS compressible fluid deformation algorithm performs the nonlinear registration instead of the FNIRT elastic deformation algorithm. A previous comparison of over 12 registration methods across multiple datasets show that ANTs consistently outperforms FNIRT on every performance metric(Ou et al., 2014). The pipeline provides an option to use a study-specific template as an intermediate image for atlas registration, which can help improve the registration in populations with systematic structural differences.

###### 4) Functional volume mapping (Vol) changes

Only one modification was made to the functional mapping (“vol” and “surf”) stages of the HCP pipeline, where functional data are projected onto the standard template and converted to CIFTI format. The unmodified HCP pipeline required the use of reverse phase spin-echo or gradient field maps in order to complete the functional mapping. Such images are used to correct distortions produced when acquiring fMRI data. While such distortion correction is considered best practice in modern fMRI studies, the modified HCP pipeline includes a “no distortion correction” approach, when no such data is acquired. Such an approach is a necessity for some large legacy datasets (e.g. ABIDE). Like most fMRI studies, the pipeline performs slice timing correction, mode 1000 normalization and frame-frame realignment at this stage (DCAN Preproc Figure 1F). Framewise displacement (FD) measures, are used in the DCAN preproc stage below to help control for the motion artifact.

#### 6) DCAN Preproc

We include a new module to the HCP pipeline, termed “DCAN Preproc”. This module performs functional connectivity preprocessing on rest fMRI scans only. Since the discovery of the motion artifact by multiple independent labs, many researchers conducted studies to determine the best standards and practices for quality assessment of functional connectivity data(Ciric et al., 2017; Power et al., 2017, 2019a; Satterthwaite et al., 2013). This module attempts to incorporate such standards here (DCAN Preproc Figure 1) and involves four broad steps: 1) standard preprocessing, 2) application of a respiratory motion filter, 3) motion censoring followed by standard reprocessing, and 4) construction of parcellated timeseries

##### *Standard preprocessing*

Standard preprocessing comprises three steps. First all fMRI data are demeaned and detrended with respect to time. Next a general linear model is used to denoise the processed fMRI data. Denoising regressors comprise signal and movement variables. Signal variables comprise mean time series for white matter, CSF, and the mean CIFTI timeseries signal; white matter and CSF are derived from Individualized segmentations generated during 3) PostFreesurfer (Figure 1C), while the mean CIFTI timeseries signal is our proxy for the global signal (see below: On global signal regression). Movement variables comprise translational (X,Y,Z) and rotational (roll, pitch, and yaw) measures estimated by realignment during *VOL* (Figure 1D) and their Volterra expansion. These regressors represent the best approach to denoising currently studied(Ciric et al., 2017). In particular, accounting for the noise regressed out by the global signal is critical for most resting-state functional MRI comparisons, as demonstrated empirically by multiple independent labs(Ciric et al., 2017; Power et al., 2017, 2019b; Satterthwaite et al., 2013). After denoising the fMRI data, the timeseries are bandpass filtered between 0.008 and 0.09 Hz using a 2$^{nd}$ order butterworth filter. Such a bandpass filter is softer than other filters, and avoids potential aliasing of the timeseries signal.

##### *On Global Signal Regression*

Though our pipeline uses the mean CIFTI timeseries instead of the global brain signal, we feel it is important to comment on the importance of global signal regression (GSR) for current functional connectivity research practices. GSR has been consistently shown to reduce the effects of motion on BOLD signals and eliminate known batch effects that directly impact group comparisons (Ciric et al., 2017; Power et al., 2015, 2019b). Motion censoring (see below) combined with GSR has been shown to be the best existing method for eliminating artifacts produced by motion. Despite multiple replications by independent labs, GSR remains a controversial procedure. GSR generally receives two criticisms, which we will address here.

First, a common critique is that GSR centers the distribution of functional correlations and produces spurious negative correlations, which do not exist without GSR. Furthermore, models of BOLD signals show that GSR distorts the correlation between regions. While it is true that GSR displays biases towards negative correlations, early(Power et al., 2015) and recent (Ciric et al., 2017) work has shown that cases where little to no motion has occurred already show negative correlations. In fact, as individuals exhibit less and less movement, functional correlations gradually shift towards the center without any GSR at all. This suggests that the "centering" effect observed corrects for some of the artifacts produced by motion. Furthermore, models that suggest GSR distorts underlying "true" correlations only used a maximum of three regions to test their hypothesis (Saad et al., 2012). As a result, the contribution of each region to the GSR was extremely high. If instead, 20 regions are used, GSR has little effect on each individual correlation, because each region contributes substantially less to the GSR; typically, functional connectivity studies use thousands of data points to compute the GSR, so one can expect little distortion.

The second common critique is that GSR can cause spurious differences when comparing populations. However, empirical studies have shown that group comparisons show fewer differences when motion and/or GSR is taken into account(Ciric et al., 2017; Power et al., 2015; Tyszka et al., 2014). If GSR is not taken into account, differences between lower and higher movers within the same population are more pronounced. Such findings strongly suggest that spurious differences are greater when GSR is not used.

Despite our strong recommendation on the importance of GSR and validating its importance, there are very specific situations where GSR is suboptimal. For example, GSR has been found to be suboptimal in macaque studies where the head is fixed to eliminate head motion and ferumoxytol is used as a contrast agent (Grayson et al., 2017). In such a case, the nature of the signal is quite different from the traditionally acquired BOLD signal, and GSR can introduce confounds into regional timeseries. Because such studies are unlikely to be conducted in human participants, our GSR proxy is turned on by default. Future versions may disable the GSR proxy if better alternatives to denoising are found.

##### *Respiratory Motion Filter*

Recent fMRI studies have started to incorporate multi-band acquisitions that enable significantly faster TRs. This enables more dense timeseries acquisitions, enabling a better estimation of the BOLD timeseries. However, such advantages come with limitations. In working with ABCD data, we have found that a respiratory artifact is produced within multi-band data (Fair et al., 2020). While this artifact occurs outside the brain, it can affect estimates of frame alignment, leading to inappropriate motion censoring (see below). Therefore, we acquire physiological data via a Physiological Monitoring Unit (PMU), and use this data to determine the respiratory signal of the participant. We then apply the respiratory signal to the FD calculated from *VOL* (Figure 1D) By filtering the frequencies of the respiratory signal from the motion realignment data, our respiratory motion filter produces better estimates of FD, which are then used for quality control.

##### *Motion censoring*

Our motion censoring procedure follows the most recent studies on eliminating the motion artifact in resting state functional MRI data (Ciric et al., 2017). Our motion censoring procedure is used for performing the standard preprocessing, and for the final construction of parcellated timeseries (Figure 1D). FD is calculated as the squared sum of all the motion vectors estimated during frame-frame alignment. For standard preprocessing, data are labeled as “bad” frames if they exceed and FD of 0.3. Such “bad” frames are removed when demeaning and detrending, and betas for the denoising are calculated using only the “good” frames. For bandpass filtering, interpolation is used initially to replace the “bad” frames, and the residuals are extracted from the denoising GLM. In such a way, standard preprocessing of the timeseries only uses the “good” data but avoids potential aliasing due to missing timepoints. When extracting timeseries for data analysis, only data with an FD of 0.2 are extracted. After motion censoring, timepoints are further censored using an outlier detection approach (Fair et al., 2020).

***Generation of parcellated timeseries for specific atlases***

Using the processed resting state fMRI data, the modified pipeline constructs parcellated timeseries for predefined atlases (Figure 1F), making it easy to construct correlation matrices or perform timeseries analysis on putative brain areas defined by independent datasets. The atlases comprise recent parcellations of brain regions that comprise different networks. In particular, parcellated timeseries are extracted for Evan Gordon’s 333 ROI atlas template (Gordon et al., 2014), Jonathan Power’s 264 ROI atlas template (Power et al., 2011), Thomas Yeo’s 118 ROI atlas template (Yeo et al., 2011), and the HCP’s 360 ROI atlas template (Glasser et al., 2016). Since we anticipate newer parcellated atlases as data acquisition, analytic techniques, and knowledge all improve, it is trivial to add new templates for this final stage.

#### 7) Executive Summary

Unlike volume-based pipelines, surface-based pipelines do not have metrics that enable simple quality control (QC; also known as quality assurance/QA) to exclude participants. Though efforts are ongoing to construct QC metrics and automated approaches to determining the quality of outputs, visual inspection of HCP outputs is critical to ensure that analyses are not contaminated by poor reconstructions, artefactual fMRI data, and improper registration. Because such outputs are visualized in multiple ways that require multiple programs, our pipeline produces an integrated, web-based executive summary, which enables visual inspection of most relevant outputs within a simple web browser. This executive summary enables one to view surfaces overlaid on MRI anatomical data (e.g. T1/T2), anatomical/functional and anatomical/anatomical overlays, and timeseries greyplots of fMRI data.

***Surfaces on T1w***

Visual inspection of surfaces on MRI volumes is time-consuming but necessary to determine whether white (i.e. defined by the grey/white border) and pial (defined by the grey/CSF border) surfaces appropriately delineate the cortical ribbon. Surfaces are overlaid on T1 weighted MRI volumetric data. Because such data are volumetric, only a single 2-dimensional slice can be visualized per image. Therefore, we converted surface/MRI overlays into a series of gif snapshots for coronal, sagittal, and axial views. Using brainsprite, we assembled these snapshots into three connected views. Through brainsprite, the user can easily cycle through volumetric slices to assess the quality of delineated pial and white surfaces. These visualizations capture the same information as found in the excellent VisualQC package (Raamana, 2019).

***Surfaces on T2w***

If a given user includes T2 weighted MRI data, additional brainsprite images will be produced for the pial and white surfaces overlaid on the T2 volume. Such visualizations can serve as a secondary check on surface quality, because the T2 contrast is better suited to assessing the quality of surfaces along inferior frontal and mesial temporal lobe cortex.

***Produce rest outlined on T1 and vice-versa to check FOV and functional to anatomical registration***

Within subject functional/anatomical registration can be inspected via anatomical/functional overlays. Specifically, selected coronal, sagittal and axial slices of anatomical MRI data are displayed with the edges of the functional data overlaid on top as a red line drawing and vice versa. Such visualizations are similar to those found in summary reports from FSL, which are used to assess the quality of within-subject functional registration.

***Produce atlas outlined on T1 and vice-versa to check atlas registration***

Similarly, atlas-based anatomical registration can be checked via anatomical/anatomical overlays. Here, selected coronal, sagittal, and axial slices of the subject's atlas transformed T1 weighted images are displayed with the edges of the atlas T1 images overlaid on top as a red line drawing and vice versa. These visualizations can help assess the quality of atlas registration.

***Produce greyplots for visualization of resting state data quality***

Identification of fMRI artifacts, such as motion induced artifact, benefit greatly from greyplot visualizations. Such visualizations were pioneered by the Schlaggar/Petersen labs at Washington University in St. Louis. The functional MRI data is plotted as an indexed plot of greyscale voxels. Each voxel is plotted along the y-axis, while each timepoint is plotted along the x-axis. The intensity of each voxel represents its value at a given timepoint. Above the graph, motion parameters, DVARS, and signal variance are plotted with respect to time. Below the graph, FD is plotted, and lines denote cutoffs for different thresholds. As a result, users can quickly scan this graph and determine the amount of high-quality fMRI data is available from each fMRI session.

***Produce concatenated resting-state greyplot, excluding any task runs***

Although we show both task and non-task greyplots for individual runs, greyplots are more critical for evaluating the quality of resting state fMRI data than task data. Therefore, we also show a concatenated greyplot across all rest runs and exclude any task-based fMRI data from this concatenation. Such a plot enables the user to quickly assess the amount of high-quality data across all resting state scans.

***Vertex-wise matrix generation***

Average whole-brain CIFTI dense connectivity (dconn) matrices were generated for ARMS-1 and ARMS-2 groups only. Using the dense timeseries (see: "Dcan Preproc"), we calculated the lag-zero pearson's correlation coefficient between every pair of grey-ordinates for every participant and wrote the output to a dconn file. We then calculated the mean dense connectivity matrix across every participant within each group. The correlation between the two matrices was calculated to measure the within-study reproducibility. Unfortunately, the subset reliability analysis was not feasible for vertex-wise matrices. Each dconn comprises ~8.1 billion data points and requires 32GB of storage, making the generation of all dconns in parallel unfeasible. Therefore, the mean dconn was calculated by producing each subject's dconn, adding the dconn to a summed dconn, and removing the subject. Per group, the summed dconn divided by the number of summed subjects produced the mean dconn. Such a procedure required a month of processing time.

***Parcellated connectivity generation and reproducibility analysis***

Parcellated connectivity matrices (pconns) are much smaller, and can be further explored to estimate within-study results reproducibility. Using the different sets of parcellated timeseries (see: "DCAN_preproc: Generation of parcellated timeseries for specific atlases"), we calculated the lag-zero pearson's correlation coefficient between every pair of parcellated regions of interest (ROIs). Per subject and parcellation scheme, this results in an ROI x ROI correlation matrix. To evaluate the results reproducibility of group average pconns, we calculated the average ARMS-1/2 pconns and computed correlation between ABCD -1 and ARMS-2.

***Communities defined from FC matrices***

Communities, the network-organization of functionally distinct brain units, were derived using the Infomap algorithm, which has been identified to work best on FC MRI data, and previously used to identify communities across whole-brain data(Gordon et al., 2014). Briefly, FC matrices were thresholded with density thresholds from 1% to 5%. Communities with fewer than three nodes were considered junk and comprised a single community. Community matrices were derived on average FC matrices for ARMS-1/2 and ABIDE-1/2 groups separately. Results reproducibility was assessed by measuring the normalized mutual information (NMI) between split-half groups. Values range from 0 to 1, with 0 indicating no overlap between community assignments and 1 indicating complete overlap.

#### ABCD-BIDS tfmri (task-fMRI) pipeline

Abcd-bids-tfmri, a modified version of the TaskfMRIAnalysis stage of the HCP-pipeline (Glasser et al., 2013) developed at University of Vermont, was used to process task-fmri data from the minimally processed ABCD-BIDS (Feczko et al., 2020b) processing pipeline (v.1.0) data, as well as derived ABCC data (Feczko, 2020; ABCD-3165). Given abcd-bids-tfmri pipeline's focus of reproducibility in neuroimaging, it allows for minimal user input while providing vast flexibility with regard to the tfMRI data that can be processed (including the type of task and the number of subject-level runs). Transparency is easily achieved with the abcd-bids-tfmri pipeline as users can efficiently share their command-line that was used in processing their data when presenting their findings.

Given its focus on dtseries data, the abcd-bids-tfmri pipeline heavily relies on HCP workbench commands (https://www.humanconnectome.org/software/workbench-command). This includes completing the user-specified spatial smoothing (wb_command -cifti-smoothing), converting the smoothed data to and from a format that FSL (Jenkinson et al. 2012) can interpret (wb_command -cifti-convert), separating the dtseries data into its comprised components (wb_command -cifti-separate-all), and reading in pertinent information from the dtseries data (wb_command -file-information), among others. Based on the user-specified parameters for censoring volumes (i.e. initial and/or high-motion frames), the pipeline will read in the filtered motion file (Fair et al., 2020) produced by the ABCD-BIDS processing pipeline and create a matrix for nuisance regression. Finally, highpass filtering is completed before running FSL's FILM (Woolrich et al. 2001).

For FILM to run, users must supply their own subject-, task-, and run-specific event timing files that are in the FSL standard three column format (i.e. onset, duration, weight/magnitude). Additionally, users need to supply a task-specific fsf template file per task that they will be processing using the abcd-bids-tfmri pipeline. As the abcd-bids-tfmri pipeline modifies this template to make it subject- and run-specific, certain values need to be replaced with specific variables that the abcd-bids-tfmri pipeline will be able to recognize. An example fsf template for ABCD's MID task is made available for users to review on ABCC (https://osf.io/psv5m/).

Users can specify which task data they would like to process by providing a list of task names within the abcd-bids-tfmri pipeline's command line interface. If the user specifies multiple runs of the task, the pipeline will complete higher-level analyses (i.e. fixed effects modeling) to combine a given subject's run-level data. Therefore, if a study has three different fMRI tasks that consist of two runs, all six level 1 analyses and all three level 2 analyses can be completed for a subject with a single run of the abcd-bids-tfmri pipeline.

The outputs of the abcd-bids-tfmri pipeline include the fully-processed dtseries data that are subsequently ready for the user to perform their desired third-level or group-wise analyses. Future releases of this pipeline will include support for volumetric/voxelwise tfmri data processing. The stable-release version of abcd-bids-tfmri is available through the ABCC (https://osf.io/psv5m/).

#### ABCD ABCC file structure

<table>
<tr><th>FILE OR FOLDER NAME</th><th>DESCRIPTION</th></tr>
<tr><td>ABCD-BIDS (where #### means sub-SUBID_ses-SESID)</td><td>Root folder of the whole ABCD-BIDS dataset</td></tr>
<tr><td>├── dataset_description.json</td><td>A JSON file describing the dataset</td></tr>
<tr><td>├── README</td><td>A free form text file describing the dataset in more detail</td></tr>
<tr><td>├── CHANGES</td><td>Version history of the dataset (describing changes, updates and corrections)</td></tr>
<tr><td>├── task-(MID|nback|rest|SST)_bold.json</td><td>BIDS-inherited common descriptions for the four major tasks</td></tr>
<tr><td>├── derivatives</td><td>A folder for data derived from processing</td></tr>
<tr><td>│ ├── freesurfer-5.3.0-HCP/sub-SUBID/ses-SESID/stats/*</td><td>FreeSurfer stats folder</td></tr>
<tr><td>│ └── abcd-hcp-pipeline</td><td>This processing is specifically the abcd-hcp-pipeline</td></tr>
<tr><td>│ ├── ##_dparc.dlabel.nii</td><td>A dense label parcellation with FreeSurfer subcorticals, where "##" is either Gordon2014FreeSurferSubcortical, HCP2016FreeSurferSubcortical, Markov2012FreeSurferSubcortical, Power2011FreeSurferSubcortical, or Yeo2011FreeSurferSubcortical corresponding to the first author and publication year</td></tr>
<tr><td>│ └── sub-SUBID</td><td>Subject-level folder</td></tr>
<tr><td>│ └── ses-SESID</td><td>Session-level folder</td></tr>
<tr><td>│ ├── img/*</td><td>DCAN Labs executive summary HTML images folder</td></tr>
<tr><td>│ ├── ####.html</td><td>DCAN Labs executive summary HTML file</td></tr>
<tr><td>│ ├── anat</td><td>Anatomical imaging derivatives</td></tr>
<tr><td>│ │ ├── ####_hemi-(L|R)_space-(MNI|T1w)_mesh-(fsLR32k|fsLR164k|native)_midthickness.surf.gii</td><td>Left or Right mid-thickness CIFTI in MNI or native T1 space with 32k, 146k, or native mesh</td></tr>
<tr><td>│ │ ├── ####_space-ACPC_dseg.nii.gz</td><td>Discrete segmentation in subject's native space in a volume</td></tr>
<tr><td>│ │ ├── ####_space-fsLR32k(_desc-smoothed)_myelinmap.dscalar.nii</td><td>Smoothed (_desc-smoothed) or unsmoothed myelin map CIFTI when a T2w image is present in the inputs</td></tr>
<tr><td>│ │ ├── ####_space-fsLR32k_(curv|sulc|thickness).dscalar.nii</td><td>Dense subject curvature, sulcal depth, or cortical thickness CIFTI</td></tr>
<tr><td>│ │ ├── ####_T1w_space-MNI_desc-wmparc_dseg.nii.gz</td><td>White matter parcellation discrete segmentation file in a volume</td></tr>
<tr><td>│ │ └── ####_(T1w|T2w)_space-MNI_(brain|head).nii.gz</td><td>Anatomical imaging masked brain or full head in MNI space in a volume</td></tr>
<tr><td>│ └── func</td><td>Functional imaging derivatives</td></tr>
<tr><td>│ ├── ####_task-(MID|nback|rest|SST)_bold_desc-filtered_timeseries.dtseries.nii</td><td>Concatenated functional task dense time series post-DCANBOLDProc (regression and filtering) in Atlas space</td></tr>
<tr><td>│ ├── ####_task-(MID|nback|rest|SST)_bold_atlas-##_desc-filtered_timeseries.ptseries.nii</td><td>Concatenated functional task parcellated time series, "atlas-##" refers to the parcellations above</td></tr>
<tr><td>│ ├── ####_task-rest_bold_atlas-###_desc-filtered_timeseries_thresh-fd0p2mm_censor-(5min|10min|belowthresh)_conndata-network_(censor.txt|connectivity.pconn.nii)</td><td>Connectivity matrix and its frame censor with either 5 minutes of data, 10 minutes of data, or all frames below the FD threshold, where "###" is Gordon2014FreeSurferSubcortical</td></tr>
<tr><td>│ ├── ####_task-(MID|nback|rest|SST)_desc-filtered(withoutliers)_motion_mask.mat</td><td>"5 contiguous frames" algorithm censoring file of temporal masks by FD threshold (0mm->0.5mm) with or without outliers</td></tr>
<tr><td>│ ├── ####_task-(MID|nback|rest|SST)_run-#_bold_timeseries.dtseries.nii</td><td>Individual functional task run dense time series in Atlas space</td></tr>
<tr><td>│ ├── ####_task-(MID|nback|rest|SST)_run-#_desc-filtered(includingFD)_motion.tsv</td><td>Movement-artifact-filtered movement numbers with or without FD</td></tr>
<tr><td>│ ├── ####_task-(MID|nback|rest|SST)_run-#(_desc-includingFD)_motion.tsv</td><td>Unfiltered raw movement numbers with or without FD</td></tr>
<tr><td>│ └── ####_task-(MID|nback|rest|SST)_run-#_space-MNI_bold.nii.gz</td><td>Motion-corrected individual functional task run in MNI space in a volume</td></tr>
<tr><td>├── sourcedata</td><td>A folder for data as acquired from the source</td></tr>
<tr><td>│ └── sub-SUBID</td><td>Subject-level folder</td></tr>
<tr><td>│ └── ses-SESID</td><td>Session-level folder</td></tr>
<tr><td>│ └── func</td><td>Functional imaging source data</td></tr>
<tr><td>│ └── ####_task-(MID|nback|SST)_run-0#_bold_EventRelatedInformation.txt</td><td>E-Prime event timings as TXT files per functional task-based run</td></tr>
<tr><td>└── sub-SUBID</td><td>Subject-level folder</td></tr>
<tr><td>└── ses-SESID</td><td>Session-level folder</td></tr>
<tr><td>├── anat/####(_rec-normalized)_(T1w|T2w).(json|nii.gz)</td><td>Each anatomical image with optional "normalized reconstruction" for SIEMENS scans</td></tr>
<tr><td>├── dwi/####_dwi.(bval|bvec|json|nii.gz)</td><td>Diffusion imaging input data</td></tr>
<tr><td>├── fmap/####(_acq-dwi)_dir-(AP|PA)_run-0#_epi.(json|nii.gz)</td><td>Each field map run pair with phase encoding direction A->P or P->A, including diffusion-specific field maps</td></tr>
<tr><td>└── func/####_task-(MID|nback|rest|SST)_run-0#_bold.(json|nii.gz)</td><td>Each functional task and each run</td></tr>
</table>

**Table S1 Detailed ABCC data structure table.** Layout of ABCD 3165 data collection. The table shows the organization of the subfolders within the 3165 dataset. The descriptions provided in the table describe each file in detail. The description here is the same for Table 1 and discusses each subsection from top to bottom. **(Yellow)** The dataset comprises a root folder describing the whole dataset. **(Orange)** Derived data can be found in the derivatives subfolder, organized by pipeline. For example, researchers that want access to our pipeline derived outputs for analysis can find them in the “abcd-hcp-pipeline” folder. **(Blue)** Researchers can find additional data, such as the event files needed for task runs, in the “sourcedata” subfolder. **(Green)** The BIDS converted input data can be found in each “sub-SUBID” directory for researchers interested in examining data processing strategies.

A more detailed organization of the ABCD ABCC dataset structure is shown in Table S1. As in Table 1, the dataset structure is depicted in sections. Researchers can check information about the dataset version and changes in the README and json files (Table S1; yellow). Derivatives used for analysis are organized by pipeline subfolder (Table S1; orange). ABCD-BIDS pipeline outputs are organized into anatomical and functional derivatives. Anatomical derivatives include surface structure and surface metric files, enabling researchers to investigate metrics like cortical thickness or myelin content. Functional derivatives are produced for each task and rest run, including post-regression dense and parcellated timeseries CIFTI files, motion files for frame censoring, and volumetric processed data. Source data (Table S1; blue) includes subject specific data produced during session acquisitions, such as event data during task scans. Finally, researchers have access to the BIDS converted input data (Table S1; green) if they want to reprocess data using different pipelines.

#### ABCD Reproducible Matched Samples (ARMS)

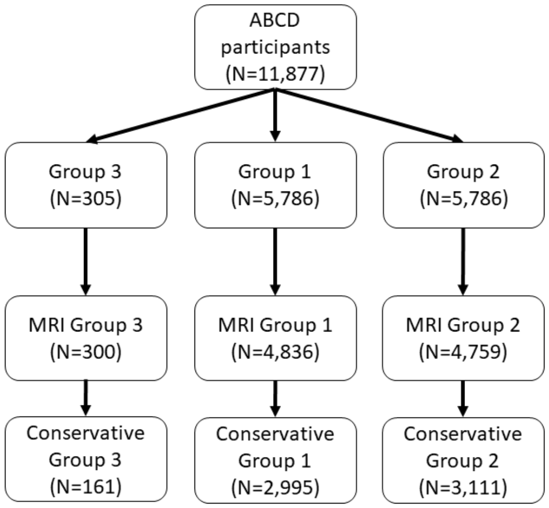


**Figure S2. The complete ABCD dataset was split into ARMS-1, ARMS-2 , and ABCD-3 stratified groups.** A flow chart depicts how ABCD data were split into three subgroups. First, three stratified groups were genereated from the entire ABCD dataset. Next, MR data from each group were processed through the DCAN-ABCD BIDS pipeline. Finally, a subset of the data, comprising participants with at least 10 minutes of clean resting state data, were selected for further analysis.

As described in the main manuscript, ABCD data (N=11,877) were stratified into three groups (Figure S2). Two larger split-halves, ARMS-1 and ARMS-2, matched on 9 demographic variables (N=5,786) and a smaller homogenous sample, ABCD-3 (N=305), used as a template (N=305).  ARMS-1 (N=4,836), ARMS-2 (N=4,759), and ABCD-3 (N=300) cases with available functional MRI data underwent MRI processing through the ABCD BIDS pipeline. After connectivity preprocessing, data quality were further examined based on head motion. Analyses proceeded with all cases with 10 minutes or greater of resting state data for ARMS-1 (N=2,995), ARMS-2 (N=3,111), and ABCD-3 (N=161). Histograms showing the demographic distributions by ARMS-1 and ARMS-2 show that the groups are highly similar on matched demographic variables (Figures S3 and S4).

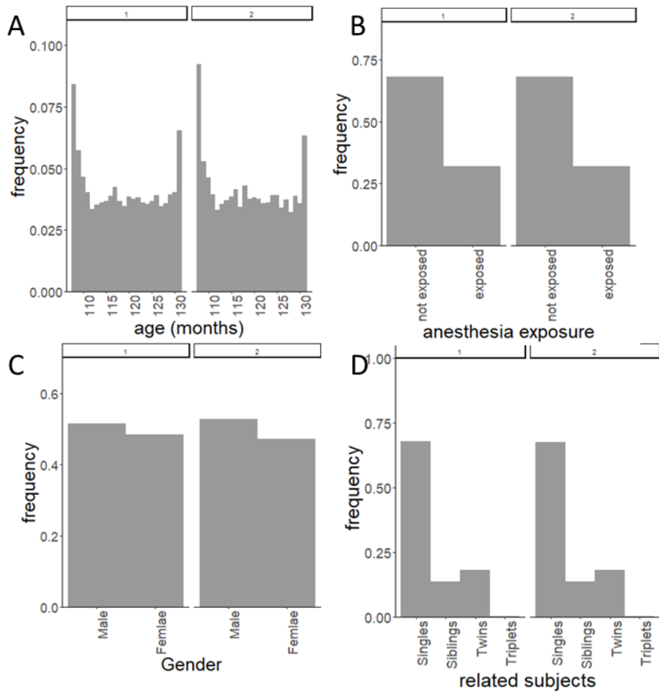


**Figure S3. Histograms of six demographic factors for ARMS-1 and ARMS-2** Distributions of six of the demographics are shown split by ARMS-1 and ARMS-2 subgroups. X-axis depicts the demographic measure, while the y-axis depicts frequency per bin. **(A)** Plot shows distribution of maximum parental education by subgroup. **(B)** Plot shows site distribution for subgroups. **(C)** Plot shows ethnicity distribution for subgroups. **(D)** Plot shows handedness distribution for subgroups. **(E)** Plot shows current grade distribution by subgroups. **(F)** Plot shows combined family income distributed by subgroups. Family income was defined as bins on the x-axis.exposure. (C) Gender. (D) Related subjects.

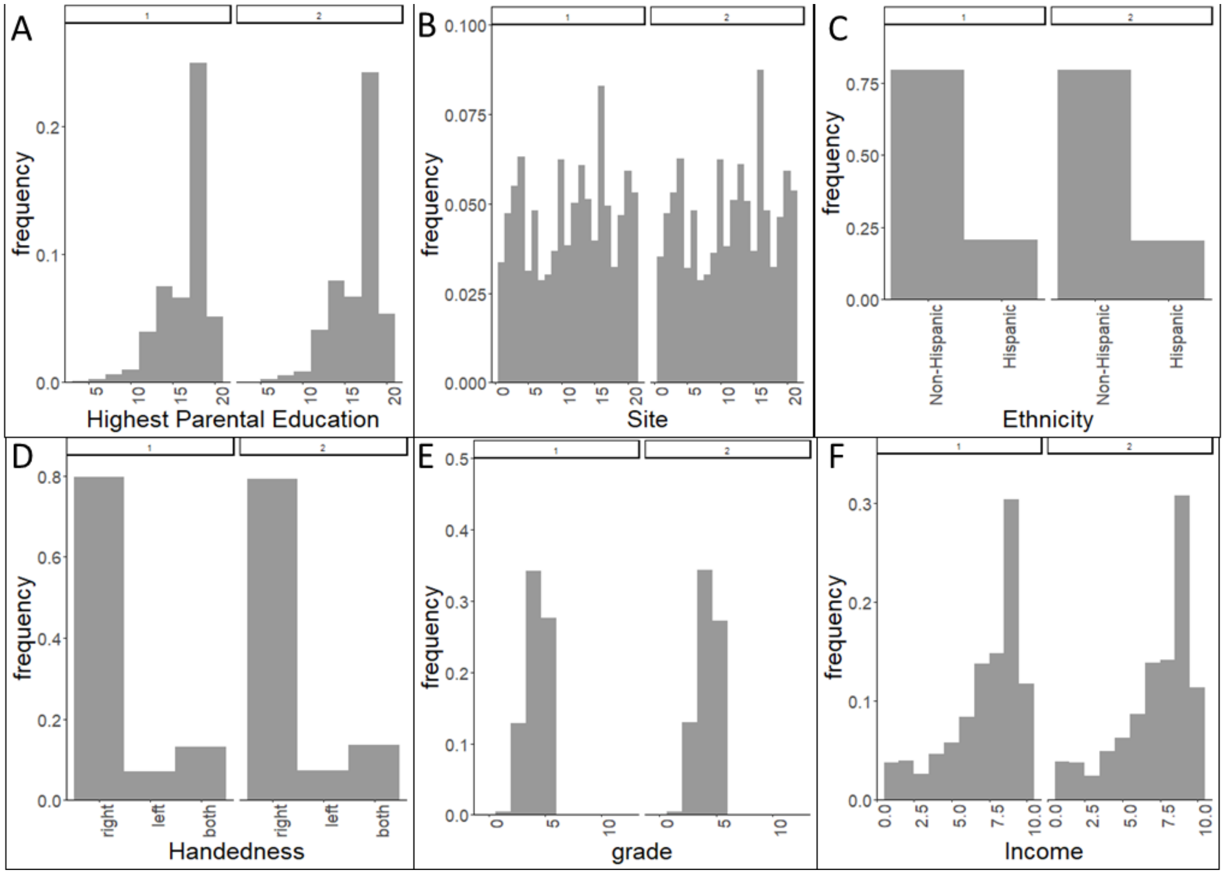


**Figure S4. Histograms of age, anesthesia exposure, sex, and sibling status demographic factors for ARMS-1 and ARMS-2.** Distributions of the remaining three demographic measures are shown, split by ARMS-1 and ARMS-2 subgroups. X-axis depicts the demographic measure, while the y-axis depicts frequency per bin. **(A)** Plot shows distribution of age by subgroup. **(B)** Plot shows anesthesia exposure for subgroups. **(C)** Plot shows sex by birth distribution for subgroups. **(D)** Plot shows sibling status distribution for subgroups.

**ABCD Resource Demographics table**

| continuous | Group1 (N=2995) mean (sd) | Group2 (N=3111) mean (sd) | Group3 (N=161) mean (sd) |
|---|---|---|---|
| age (months) | 119.64 (7.48) | 119.75 (7.47) | 118.37 (7.73) |
| demo_ed_v2 | 4.27 (0.78) | 4.27 (0.78) | 4.20 (0.76) |
| max parent ed. | 17.38 (2.85) | 17.34 (2.46) | 16.83 (2.90) |
| combined inc. | 7.51 (2.24) | 7.46 (2.24) | 7.08 (2.35) |
| **categorical** | **Group1 (N=2995) count (%)** | **Group2 (N=3111) count (%)** | **Group3 (N=161) count (%)** |
| # female | 1411 (47.10) | 1544 (49.66) | 78 (48.45) |
| # anesthesia | 966 (32.2) | 1005 (32.3) | 42 (26.1) |
| # right handed | 2401 (80.2) | 2525 (81.2) | 136 (84.5) |
| **race** | **Group1 (N=2995) count (%)** | **Group2 (N=3111) count (%)** | **Group3 (N=161) count (%)** |
| white | 2399 (80.10) | 2460 (79.07) | 106 (65.84) |
| black | 539 (18.00) | 556 (17.87) | 30 (18.63) |
| AIAK | 94 (3.14) | 99 (3.18) | 4 (2.48) |
| NHPI | 19 (0.63) | 18 (0.58) | 4 (2.48) |
| asian | 173 (5.78) | 198 (6.36) | 9 (5.59) |
| other | 143 (4.77) | 166 (5.34) | 21 (13.04) |
| unkown | 23 (0.77) | 34 (1.09) | 4 (2.48) |
| combined | 2970 (99.17) | 3070 (98.68) | 97 (60.25) |
| latinx | 544 (18.16) | 564 (18.13) | 28 (17.39) |

| site | Group1 (N=2995) count (%) | Group2 (N=3111) count (%) | Group3 (N=161) count (%) |
|---|---|---|---|
| 1 | 41 (1.37) | 49 (1.58) | 4 (2.48) |
| 2 | 212 (7.08) | 205 (6.59) | 9 (5.59) |
| 3 | 199 (6.64) | 213 (6.85) | 5 (3.11) |
| 4 | 205 (6.84) | 182 (5.85) | 9 (5.59) |
| 5 | 103 (3.44) | 112 (3.60) | 12 (7.45) |
| 6 | 170 (5.68) | 170 (5.46) | 22 (13.66) |
| 7 | 91 (3.04) | 84 (2.70) | 4 (2.48) |
| 8 | 62 (2.07) | 88 (2.83) | 3 (1.86) |
| 9 | 122 (4.07) | 112 (3.60) | 6 (3.73) |
| 10 | 128 (4.27) | 146 (4.69) | 5 (3.11) |
| 11 | 133 (4.44) | 122 (3.92) | 4 (2.48) |
| 12 | 42 (1.40) | 57 (1.83) | 3 (1.86) |
| 13 | 157 (5.24) | 145 (4.66) | 5 (3.11) |
| 14 | 189 (6.31) | 194 (6.24) | 11 (6.83) |
| 15 | 89 (2.97) | 95 (3.05) | 8 (4.97) |
| 16 | 380 (12.69) | 404 (12.99) | 10 (6.21) |
| 17 | 97 (3.24) | 121 (3.89) | 8 (4.97) |
| 18 | 88 (2.94) | 86 (2.76) | 2 (1.24) |
| 19 | 124 (4.14) | 115 (3.70) | 12 (7.45) |
| 20 | 183 (6.11) | 212 (6.81) | 10 (6.21) |
| 21 | 180 (6.01) | 199 (6.40) | 9 (5.59) |

**Table S2. Demographics for imaging data remain highly similar.** Demographics for the MRI datasets are depicted in this table. As shown, differences between ARMS-1 and ARMS-2 are fairly minimal, with gender showing the largest difference at 2.5 percent. No other factor shows differences greater than 1.03 percent. ABCD-3 differences are larger for both site and race but not other demographic factors.

Demographics for the imaging dataset are shown in Table S2. Despite the data trimming, ARMS-1 and ARMS-2 are similarly matched on demographic variables. Gender shows the largest difference with 2.5 percent more males in ARMS-2 than ARMS-1. No other variable shows more than a 1.03 percent difference between the two larger subsets. As noted in the main manuscript, ABCD-3 is more homogenous and not used for the reproducibility and stability analyses.

#### Marginal model framework

##### Marginal model details

The framework for MarginalModelCifti is similar to work by Tom Nichols and Bryan Guillaume, and detailed extensively there(Guillaume et al., 2014). Here, we briefly summarize the framework here and depict it in Figure S5.

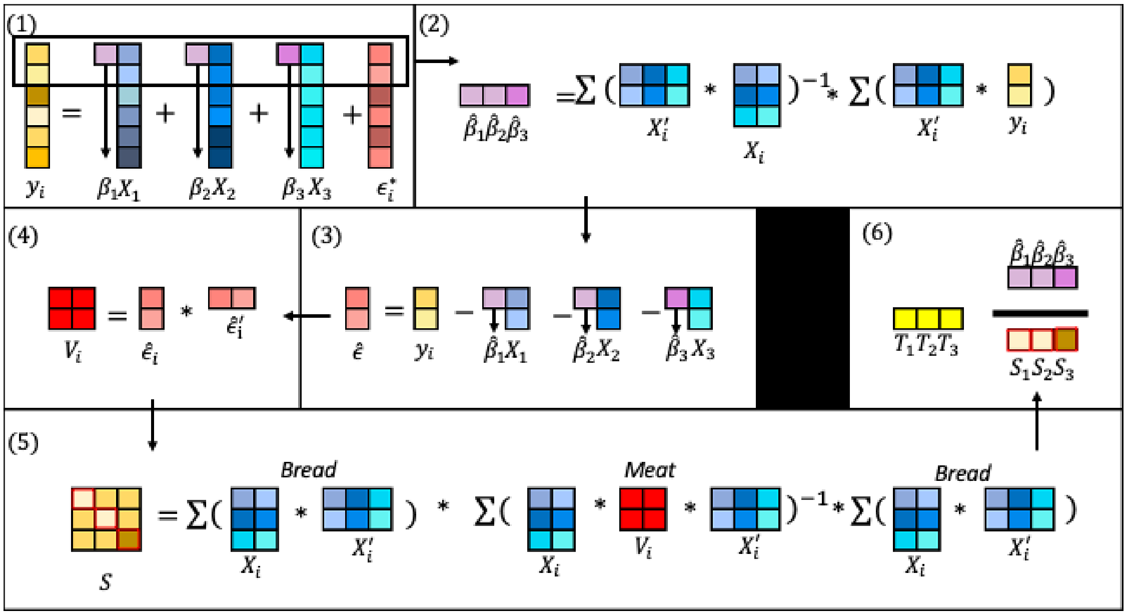


**Figure S5. Graphical representation of Marginal Model Framework.** Subfigures are noted by the headings. (1) The marginal model. Rows represent n observations of individuals and columns represent multiple predictors with betas. Each beta is a scalar associated with each column of $x$, as shown by the arrows. (2) The estimator for the betas in the marginal model for a single connection/grey-ordinate, as indicated by the black rectangle in (1). Rows represent observations and columns represent multiple predictors for individuals for group *i*. Betas are calculated by summing over all groups *m*. (3) The residuals estimated from the marginal model betas. Rows represent observations and columns represent multiple predictors or betas. Each beta is a scalar associated with each column of $x$, as shown by the arrows. (4). The covariance matrix of the residuals. Rows and columns represent observations. (5) The sandwich estimator, where the top portion represents the “meat” and the bottom two portions represent the “bread”. Rows and columns are consistent with prior definitions for each variable. (6) The calculation of the test statistic. Columns represent multiple betas.

Similar to linear mixed effects models, marginal models are linear models of the form:

$$y_i = X_i\beta + \epsilon_i^* \quad (1)$$

$y_i$ is the vector of n observations for each individual within a nested group $m$. $X_i$ is the design matrix for *n* observations by *p* fixed effects. β is the vector of p fixed effects model parameter estimates. Therefore, β can be estimated via:

$$\hat{\beta} = \left(\sum_{i=1}^{m} X_i'X_i\right)^{-1}\left(\sum_{i=1}^{m} X_i'y_i\right) \quad (2)$$

Though other forms can be used to estimate fixed effects parameters, such forms require iterative solutions that are much slower while achieving only minimal gains(Guillaume et al., 2014).

$\epsilon_i^*$ in equation (1) are the individual errors about the marginal model with mean of 0 and covariance of $V_i$. For the marginal model, the covariance matrix, $\hat{V}_i$ can be estimated via the residuals :

$$\hat{\epsilon}_i = y_i - X_i\hat{\beta} \quad (3)$$

$$\hat{V}_i = \hat{\epsilon}_i\hat{\epsilon}_i' \quad (4)$$

The covariance matrix of $\hat{\beta}$ can be estimated using the sandwich estimator via the formula:

$$S = \left(\sum_{i=1}^{m} X_i X_i\right)^{-1} \left(\sum_{i=1}^{m} X_i \hat{V}_i X_i\right) \left(\sum_{i=1}^{m} X_i X_i\right)^{-1} \quad (5)$$

Here, $\left(\sum_{i=1}^{m} X_i X_i\right)^{-1}$ is the "bread" and $\left(\sum_{i=1}^{m} X_i \hat{V}_i X_i\right)$ is the "meat" giving the sandwich estimator its name. Test statistics per parameter $p$ can be computed via:

$$T_p = \hat{\beta}_p / S_{pp} \quad (6)$$

$T_p$ is a wald-statistic, which can then be converted to z-scores for subsequent analysis.

#### Enrichment analysis

On the resulting z-score matrix, we performed an enrichment analysis (Eggebrecht et al., 2017) using the wild bootstrap in place of a permutation test. An enrichment analysis enables us to test behavior-brain associations at the level of systems and not just ROIs. Per bootstrap, a null dataset was generated and mass-univariate marginal models were fitted as described above. Z-score matrices were then thresholded for "significant" values, where z-scores greater than 2.36 were considered significant; this threshold was selected *a priori* and is consistent with prior imaging literature(Eklund et al., 2016). Using the defined community structure for the Gordon ROIs, we calculated a chi-squared statistic for every module, representing either between or within system connections. Repeating this procedure over 1000 bootstraps produces a null distribution of chi-squared statistics per module. We then compare the observed chi-squared statistic to the null distribution to calculate the p value.

#### Experiment I

##### ARMS Subset Generation for Reproducibility and Reliability Analyses

To help address questions of reproducibility, the ABCC provides researchers with a tool that extends ARMS, called ARMS Reproducibility Tool (ARMS-RT). ARMS-RT enables researchers to estimate the within-study reproducibility and reliability. ARMS-RT generates a series of subsets from one group (e.g., ARMS-1) that is demographically matched to the other ARMS (e.g. ARMS-2). First, the approach generates random subsets of one group at every sample size. A series of chi-squared tests are used to test whether each demographic factor significantly differs between each subset and the larger group. Once we identify subsets that show no statistical differences in demographics, we used the 10 demographic factors used for ARMS to calculate the Euclidean distance between the matched subset and the larger group for a given sample size. This Euclidean distance becomes the threshold for subset selection. We then generated 100 subsets where the Euclidean distance between the subset and the larger group do not exceed the calculated threshold. Sample sizes less than 25 cannot be matched on sibling relationships, but are matched on the 9 other demographic factors.

ARMS-RT does not depend on ABCD or ABCC data, and can be run on any independent dataset as well. For example, a researcher could use ARMS-RT to identify matched subsets from the UK BioBank, or use ARMS-RT to identify subsets of their own data matched to ABCD. The matched subsets provided by ARMS-RT enables researchers to estimate the sample size needed to achieve **reproducibility** for measurements like parcellated connectivity or statistical tests such as those conducted via MarginalModelCifti (see below). One can also estimate the **reliability** of effect sizes, such as cohen's d. Examples of both are shown in Section III to highlight cautionary notes when conducting mass-univariate statistical tests.

##### Connectivity Matrix Generation:

The ABCD-BIDS pipelines produce timeseries for several commonly used parcellations for connectivity matrix generation (see: "DCAN_preproc: Generation of parcellated timeseries for specific atlases" in supplemental materials). For the current examples, we calculated the lag-zero pearson's correlation coefficient between every pair of parcellated regions of interest (ROIs) for the Gordon parcellation (citation) using the CiftiConn tool (Git page here). Correlation matrices were generated for varying amounts of 'good, movement free' data. These included both 5 and 10 minute trimmed datasets per subject. This manipulation allows reproducibility comparisons as a function of the amount of data utilized per subject.

#### Experiment II

We followed the same procedure published previously for the BPPCA for ARMS-1 and ARMS-2 separately (Thompson et al., 2019). Briefly, three cognitive traits were extracted from measures from nine neurocognitive tasks using BPPCA (see supplemental materials for more details). A prior study describes the measures and their justification (Luciana et al., 2018). The first three principal component scores were derived per participant. These components best represent three cognitive traits: general cognitive ability (PC1), executive function (PC2), and learning/memory (PC3). Comparisons of cognitive trait scores show similar distributions for ARMS-1 and ARMS-2 for general cognitive ability (Figure S6A), executive function (Figure S6B), and learning/memory (Figure S6C). These comparisons mirror the distributions found in the full dataset, suggesting that the PCs here are highly reproducible for the full ARMS-1 and ARMS-2 dataset.

##### Cognitive traits extracted from ARMS are reproducible

A Bayesian probabilistic principal components analysis (BPPCA) was used to extract cognitive traits from ARMS-1 and ARMS-2 separately. We followed the same procedure published previously(Thompson et al., 2019). Briefly, three cognitive traits were extracted from measures from nine neurocognitive tasks using BPPCA. A prior study describes the measures and their justification(Luciana et al., 2018). Seven measures came from the NIH cognitive toolbox. The Toolbox Picture Vocabulary Task measures verbal ability(Gershon et al., 2013). The Toolbox Flanker Task measures the effect of context on visual stimulus responses(Eriksen and Eriksen, 1974). The Toolbox List Sorting Working Memory Test measures working memory by asking participants to sequence a series of stimuli by varying characteristics(Tulsky et al., 2013). The Toolbox Dimensional Change Card Sort Task measures the ability to maintain and switch task instructions(Zelazo et al., 2013). The Toolbox Pattern Comparison Processing Speed Test measures rapid visual processing (Carlozzi et al., 2013). The Toolbox Picture Sequence Memory Test measures an individual’s memory of sequences of actions(Bauer et al., 2013). The Toolbox Oral Reading Recognition task measures an individual’s ability to pronounce single words(Gershon et al., 2013). Measures from two additional tasks were also included. The Rey Auditory Verbal Learning Test measures auditory learning ability(Daniel et al., 2014). The little man task measures mental rotation ability(Acker and Acker, 1982). The first three principal component scores were derived per participant. These components best represent three cognitive traits: general cognitive ability (PC1), executive function (PC2), and learning/memory (PC3).

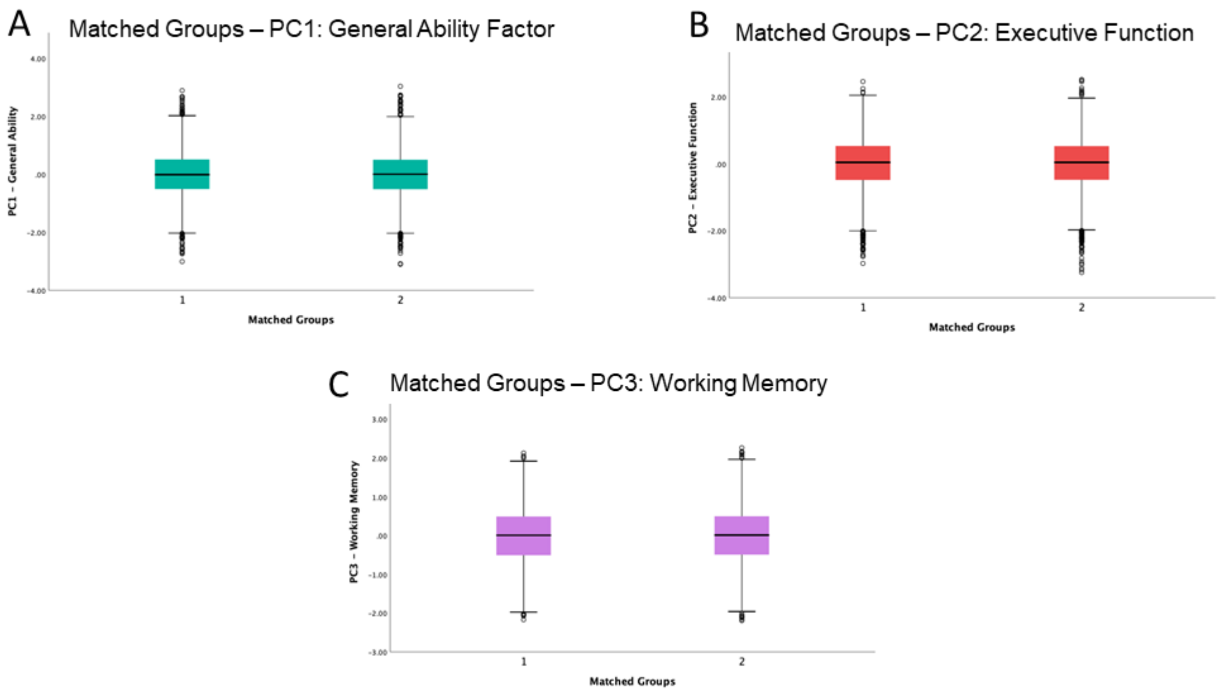


**Figure S6. Comparisons of cognitive traits show no differences between ARMS-1 and ARMS-2**. Boxplots for each cognitive trait are shown. The X-axis depicts the subgroup (ARMS-1; left, ARMS-2; right). The Y-axis depicts the trait score. **(A)** Boxplot of general ability factor for the subgroups. **(B)** Boxplot of executive function factor for the subgroups. **(C)** Boxplot of learning/memory factor for the subgroups.

Comparisons of cognitive trait scores show similar distributions for ARMS-1 and ARMS-2 for general cognitive ability (Figure S6A), executive function (Figure S6B), and learning/memory (Figure S6C). These comparisons mirror the distributions found in the full dataset, suggesting that the PCs here are highly reproducible for the full ARMS-1 and ARMS-2 dataset.

### Results

#### Experiment I: Within study reproducibility, subset reliability, and subset stability for parcellated connectivity data using other motion thresholds

##### Subset reliability and stability analysis for 5 minute trimmed datasets

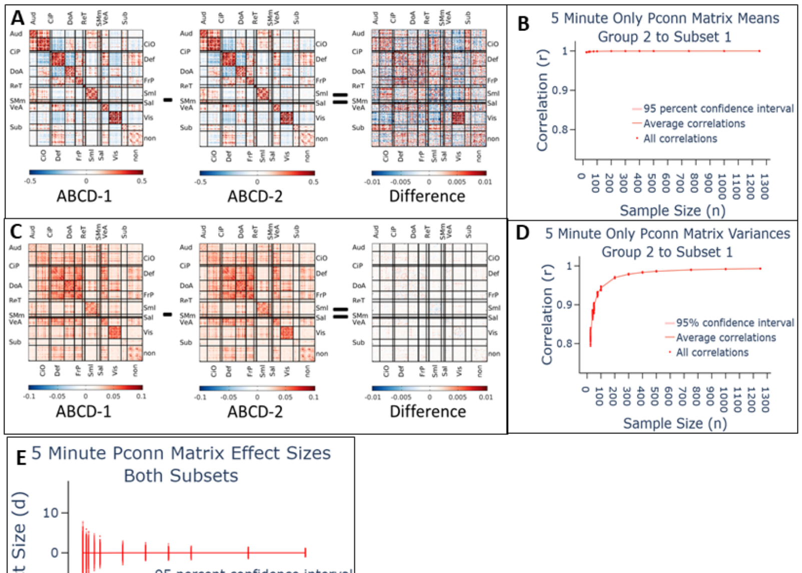


**Figure S7. Replicability of Gordon parcellated connectivity matrices from ARMS-1 and ARMS-2 datasets for 5 minutes of trimmed data. (A)** Mean connectivity matrix for ARMS-1 (left) and ARMS-2 (middle) and the difference between them (right). Correlations are sorted by the Gordon-identified community structure. The proportion of variance explained is indicated by the arrow in between the two group matrices**.** **(B)** Subset reliability analyses for ARMS-2 subsets to ARMS-1 for mean connectivity matrices. X-axis represents the sample size while Y shows the correlation. **(C)** Variance connectivity matrix for ARMS-1 (left) and ARMS-2 (middle) and the difference between them (right). Correlations are sorted by the Gordon-identified community structure. The proportion of variance explained is indicated by the arrow in between the two group matrices. **(D)** Subset reliability analyses for ARMS-2 subsets to ARMS-1 for variance connectivity matrices. X-axis represents the sample size while Y shows the correlation. **(E)** The subset stability analysis for ARMS-2 subsets to ARMS-1 for 5 minutes of data. X-axis depicts the sample size while Y shows the effect size per connection per subset.

If subjects are frame censored to precisely 5 minutes, parcellated connectivity mean matrices from ARMS-1 and ARMS-2 show high within-study reproducibility. Point estimates for ARMS-1 (Figure S7A; left) and ARMS-2 (Figure S7A; middle) were extremely similar for 5 minutes data ($r^2$=0.9992), however extremely small differences exist between the matrices (Figure S7A; right), unlike for 10 minute trimmed datasets. Subset reliability began to reach the maximum with as little as 20 subjects (Figure S7B). Parcellated connectivity variance matrices ARMS-1 (Figure S7C; left) and ARMS-2 (Figure S7D; middle), were also near identical ($r^2$=0.9913; Figure S7C; right), but sample sizes over 1000 were needed to achieve the same level of reliability (Figure S7D). Because the subset stability analyses produces stratified matched subsets, we anticipate that the "true" effect size of group subset comparisons are zero. The effect size subset stability analysis shows that observed effect sizes grow closer to zero as sample size increases (Figure S7E); however, even with 1000 participants, "true" effect sizes may still be misestimated as "large" effect size (e.g. d > 0.8).

#### Subset reliability and stability analysis for 10 minute untrimmed datasets

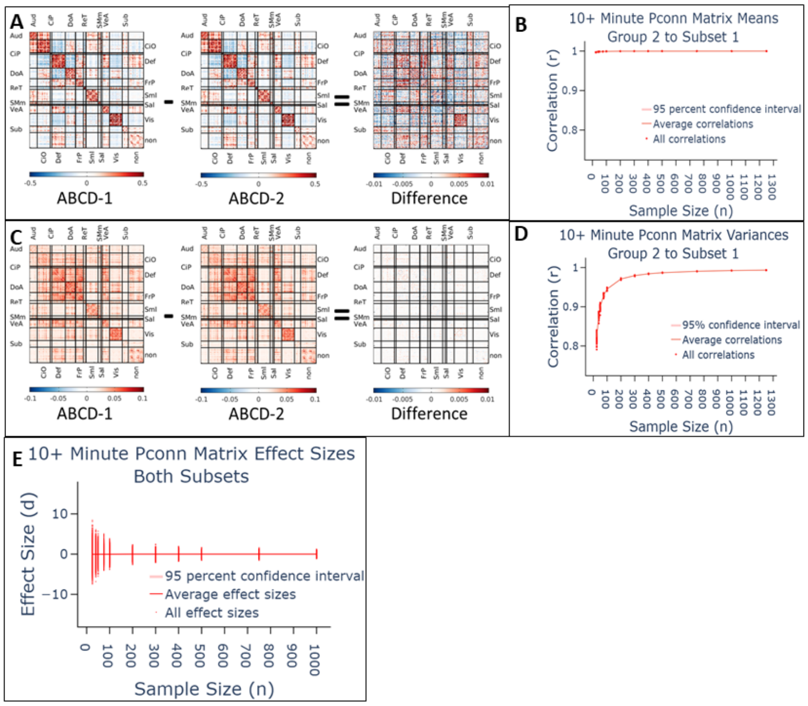


**Figure S8. Replicability of Gordon parcellated connectivity matrices from ARMS-1 and ARMS-2 datasets for 10 minutes or more of data. (A)** Mean connectivity matrix for ARMS-1 (left) and ARMS-2 (middle) and the difference between them (right). Correlations are sorted by the Gordon-identified community structure. The proportion of variance explained is indicated by the arrow in between the two group matrices. **(B)** Subset reliability analyses for ARMS-2 subsets to ARMS-1 for mean connectivity matrices. X-axis represents the sample size while Y shows the correlation. **(C)** Variance connectivity matrix for

ARMS-1 (left) and ARMS-2 (middle) and the difference between them (right). Correlations are sorted by the Gordon-identified community structure. The proportion of variance explained is indicated by the arrow in between the two group matrices. **(D)** Subset reliability analyses for ARMS-2 subsets to ARMS-1 for variance connectivity matrices. X-axis represents the sample size while Y shows the correlation. **(E)** The subset stability analysis for ARMS-2 subsets to ARMS-1 for 10 minutes of data. X-axis depicts the sample size while Y shows the effect size per connection per subset.

If subjects with more than 10 minutes are not trimmed, parcellated connectivity mean matrices from ARMS-1 and ARMS-2 show high within-study reproducibility. Point estimates for ARMS-1 (Figure S8A; left) and ARMS-2 (Figure S8A; middle) were extremely similar for 5 minutes data ($r^2$=0.9993), however extremely small differences exist between the matrices (Figure S8A; right), unlike for 10 minute trimmed datasets. Subset reliability began to reach the maximum with as little as 20 subjects (Figure S8B). Parcellated connectivity variance matrices ARMS-1 (Figure S8C; left) and ARMS-2 (Figure S8C; middle), were also near identical ($r^2$=0.9925; Figure S8C; right), but sample sizes over 1000 were needed to achieve the same level of reliability (Figure S8D). Because the subset stability analyses produces stratified matched subsets, we anticipate that the "true" effect size of group subset comparisons are zero. The effect size subset stability analysis shows that observed effect sizes grow closer to zero as sample size increases (Figure S8E); as with prior analysis, "true" effect sizes may still be grossly misestimated.

#### Subset stability analysis for 5 minute trimmed datasets

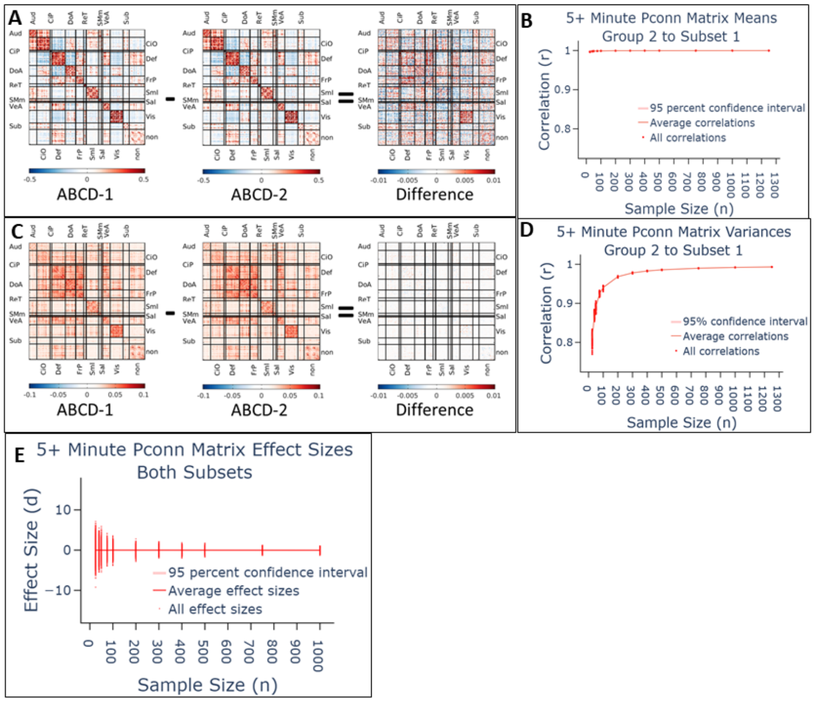


**Figure S9. Replicability of Gordon parcellated connectivity matrices from ARMS-1 and ARMS-2 datasets for 5 minutes or more of data. (A)** Mean connectivity matrix for ARMS-1 (left) and ARMS-2 (middle) and the difference between them (right). Correlations are sorted by the Gordon-identified community structure. The proportion of variance explained is indicated by the arrow in between the two group matrices. **(B)** Subset reliability analyses for ARMS-2 subsets to ARMS-1 for mean connectivity matrices. X-axis represents the sample size while Y shows the correlation. **(C)** Variance connectivity matrix for ARMS-1 (left) and ARMS-2 (middle) and the difference between them (right). Correlations are sorted by the Gordon-identified community structure. The proportion of variance explained is indicated by the arrow in between the two group matrices. **(D)** Subset reliability analyses for

ARMS-2 subsets to ARMS-1 for variance connectivity matrices. X-axis represents the sample size while Y shows the correlation. **(E)** The subset stability analysis for ARMS-2 subsets to ARMS-1 subsets for 5 minutes of data. X-axis depicts the sample size while Y shows the effect size per connection per subset.

If subjects with more than 5 minutes are not trimmed, parcellated connectivity mean matrices from ARMS-1 and ARMS-2 show high within-study reproducibility. Point estimates for ARMS-1 (Figure S9A; left) and ARMS-2 (Figure S9A; middle) were extremely similar for 5 minutes data ($r^2$=0.9994), however extremely small differences exist between the matrices (Figure S9A; right), unlike for 10 minute trimmed datasets. Subset reliability began to reach the maximum with as little as 20 subjects (Figure S9B). Parcellated connectivity variance matrices ARMS-1 (Figure S9C; left) and ARMS-2 (Figure S9C; middle), were also near identical ($r^2$=0.9931; Figure S9C; right), but sample sizes over 1000 were needed to achieve the same level of reliability (Figure S9D). Because the subset stability analyses produces stratified matched subsets, we anticipate that the "true" effect size of group subset comparisons are zero. The effect size subset stability analysis shows that observed effect sizes grow closer to zero as sample size increases (Figure S9E); however, even with 1000 participants, effect sizes may still be grossly misestimated for individual connections.

**Subset reliability analyses for ARMS-1 subsets to ARMS-2**

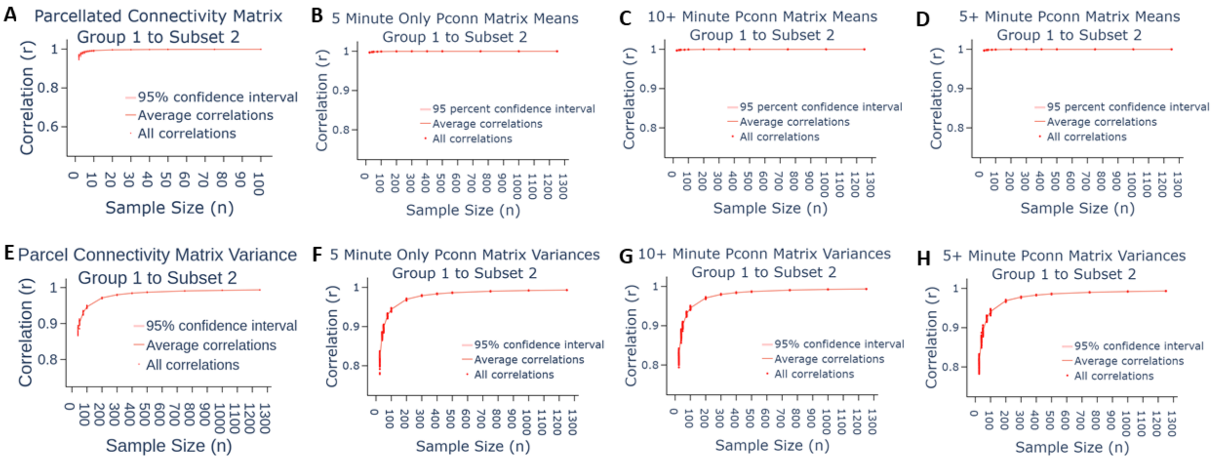


**Figure S10. Remaining subset analyses replicate findings from ARMS-1 subsets to the full ARMS-2 dataset.** All figures depict the correlation from ARMS-1 subsets to ARMS-2 dataset. The x-axis represent the given subset's sample size, while the y-axis represents the correlation between the given subset and the 10 minute ARMS-2 dataset. Subset reliability for **(A)** mean parcellated connectivity with 10 minute trimmed datasets; **(B)** mean parcellated connectivity with 5 minute trimmed datasets; **(C)** mean parcellated connectivity with 10 minute plus datasets; **(D)** mean parcellated connectivity with 5 minute plus datasets; **(E)** variance parcellated connectivity with 10 minute trimmed datasets; **(F)** variance parcellated connectivity with 5 minute trimmed datasets; **(G)** variance parcellated connectivity with 10 minute plus datasets; **(H)** variance parcellated connectivity with 5 minute plus datasets.

The manuscript so far has only discussed ARMS-2 subset to ARMS-1 reliability analyses to make the manuscript easier to read. Some may be concerned if ARMS-1 subsets to ARMS-2 show different degrees of reliability. Here, ARMS-1 subset to ARMS-2 reliability analyses are shown to illustrate that the findings are the same and alleviate this concern (Figure S10). Subset reliability analyses for 10 minute trimmed mean (Figure S10A) and variance (Figure S10E) maps replicate the findings in the main manuscript. Near perfect reliability is achieved for mean maps with as few as 20 subjects, but variance maps require more than 1000 participants to achieve the same reliability. The same findings are found for 5 minute trimmed mean (Figure S10B) and variance (Figure S10F) matrices, 10 minute untrimmed mean (Figure S10C) and variance (Figure S10G) matrices, and 5 minute untrimmed mean (Figure S10D) and variance (Figure S10H) matrices. Notably, 10 minute trimmed variance shows better reliability at small sample sizes achieves high reliability faster than the other datasets.

#### Experiment I: Within study reproducibility, subset reliability and stability analyses on curvature and sulcal depth

##### Curvature maps show high reproducibility, reliability and stability at large sample sizes

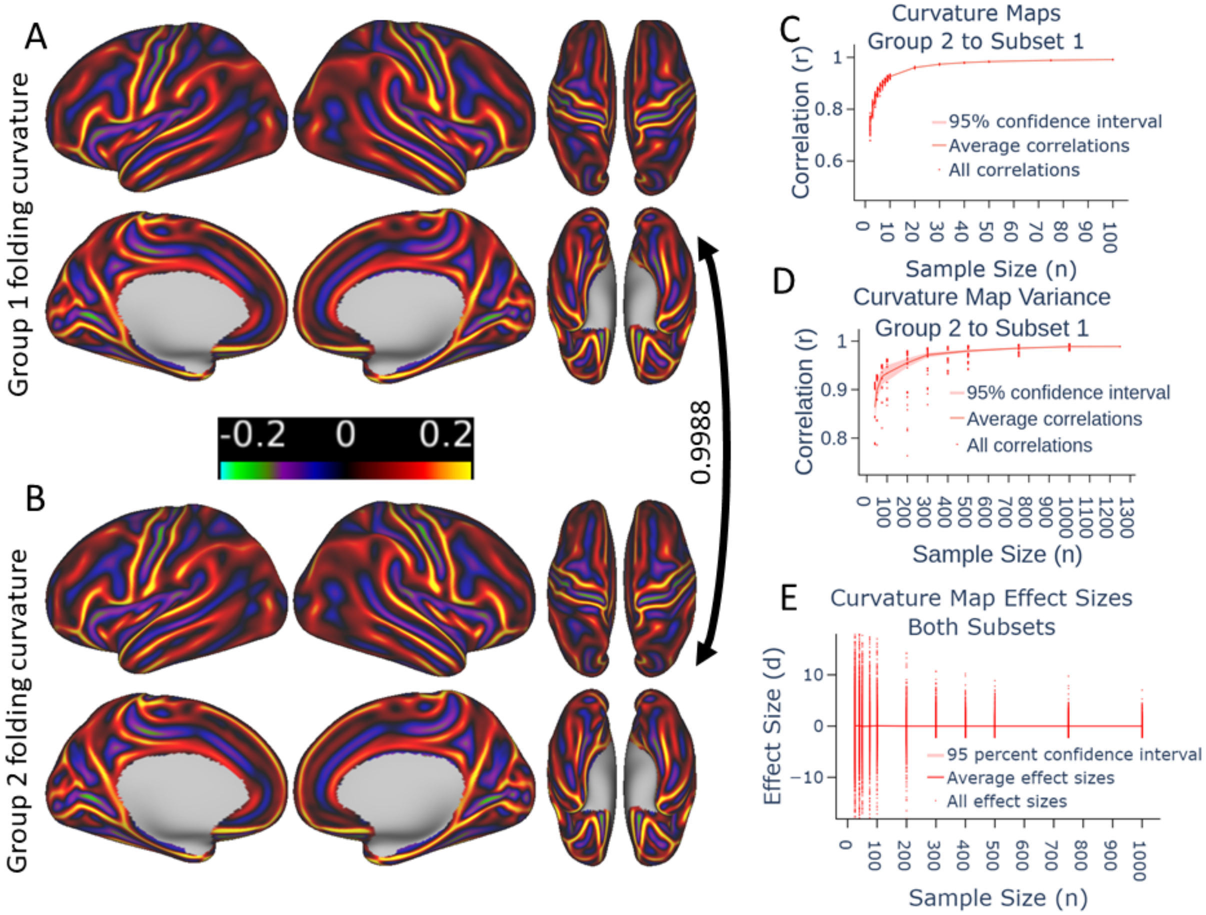


**Figure S11. Replicability of folding curvature from ARMS-1 and ARMS-2 datasets. (A)** Visualization of mean folding curvature for ARMS-1 on cortical surfaces. **(B)** Visualization of mean folding curvature for ARMS-2 on cortical surfaces. Arrow between ARMS-1 and ARMS-2 denotes the proportion of variance explained between the two maps. **(C)** Subset reliability analyses for ARMS-2 subsets to ARMS-1 for folding curvature variance. X-axis represents the sample size while Y shows the correlation. **(D)** Subset reliability analyses for ARMS-2 subsets to ARMS-1 for folding curvature mean. X-axis represents the sample size while Y shows the correlation. **(E)** Subset stability analysis for ARMS-2 subsets to ARMS-1 for folding curvature. X-axis represents the sample size while Y shows the effect size.

ARMS-1 and ARMS-2 folding curvature maps were nearly identical (Figure S11A,S11B: $r^2$=0.9992), and subset reliability of point-estimates were reliable with sample sizes as small as 100 (Figure S11C). However, variance maps required sample sizes over 1000 to achieve similar reliability (Figure S11D). Subset stability analyses show similar effect size findings as for functional connectivity analyses, where effect sizes grow closer to zero as sample size increases (Figure S11E); however, single vertex effect sizes may be grossly misestimated as “massive” effects (d > 2) when no “true” effect is present. Taken together, these results indicate that folding curvature measures are highly reliable for the ABCD dataset, but require over a thousand participants to achieve high stability.

##### Sulcal depth maps show high reproducibility, reliability, and stability at large sample sizes

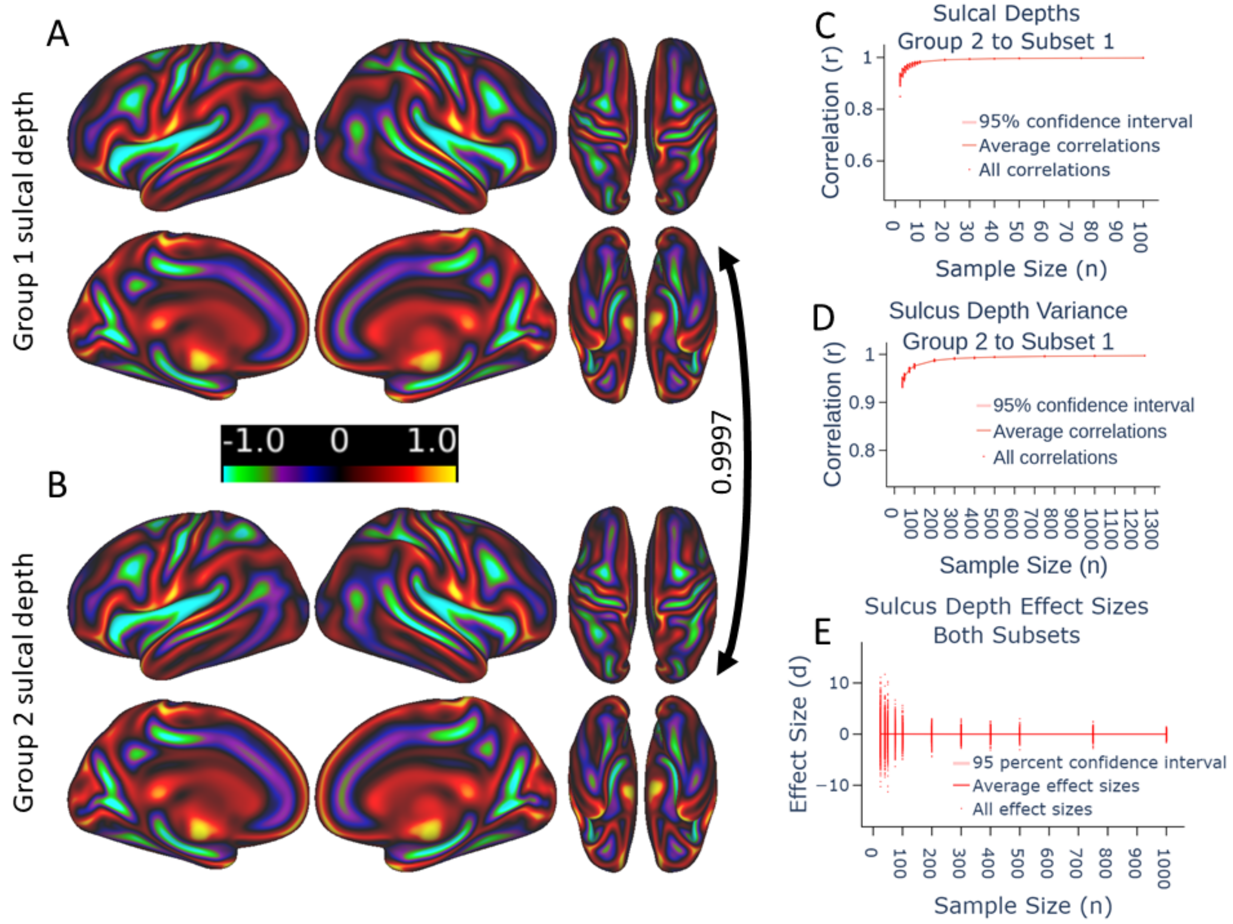


**Figure S12. Replicability of sulcal depth from ARMS-1 and ARMS-2 datasets. (A)** Visualization of mean sulcal depth for ARMS-1 on cortical surfaces. **(B)** Visualization of mean sulcal depth for ARMS-2 on cortical surfaces. Arrow between ARMS-1 and ARMS-2 denotes the proportion of variance explained between the two maps. **(C)** Subset reliability analyses for

ARMS-2 subsets to ARMS-1 for sulcal depth variance. X-axis represents the sample size while Y shows the correlation. **(D)** Subset reliability analyses for ARMS-2 subsets to ARMS-1 for sulcal depth mean. X-axis represents the sample size while Y shows the correlation. **(E)** Subset stability analysis for ARMS-2 subsets to ARMS-1 for sulcal depth. X-axis represents the sample size while Y shows the effect size.

ARMS-1 and ARMS-2 sulcal depth maps were nearly identical (Figure S12A,S12B: $r^2$=0.9992), and subset reliability of point-estimates were reliable with sample sizes as small as 100 (Figure S12C). However, variance maps required sample sizes over 1000 to achieve similar reliability (Figure S12D). Subset stability analyses show similar effect size findings as for functional connectivity analyses, where effect sizes grow closer to zero as sample size increases (Figure S12E); however, even with 1000 participants, point effect sizes could still be largely misestimated (d > 0.8). Taken together, these results indicate that sulcal depth measures are highly reliable for the ABCD dataset, but require at least 1000 subject to achieve stability for group comparisons.

#### Subset reliability analyses for ARMS-1 subsets to ARMS-2

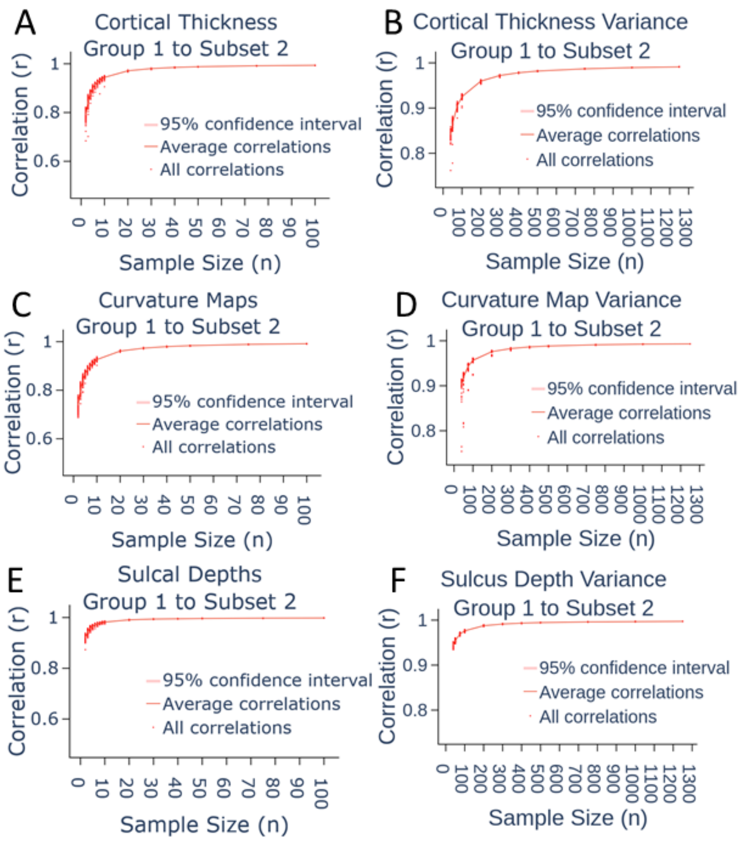


**Figure S13. Remaining subset analyses replicate findings from ARMS-1 subsets to the full ARMS-2 dataset.** All figures depict the correlation from ARMS-1 subsets to ARMS-2 dataset. The x-axis represent the given subset's sample size, while the y-axis represents the correlation between the given subset and the 10 minute ARMS-2 dataset. **(A)** Reliability for mean cortical thickness **(B)** Reliability for cortical thickness variance. **(C)** Reliability for folding curvature mean. **(D)** Reliability for folding curvature variance. **(E)** Reliability for sulcal depth mean. **(F)** Reliability for sulcal depth variance.

The manuscript so far has only discussed ARMS-2 subset to ARMS-1 reliability analyses to make the manuscript easier to read. Some may be concerned if ARMS-1 subsets to ARMS-2 show different degrees of reliability. Here, ARMS-1 subset to ARMS-2 reliability analyses are shown to illustrate that the findings are the same and alleviate this concern (Figure S13). Subset reliability analyses for cortical thickness mean (Figure S13A) and variance (Figure S13B) maps replicate the findings in the main manuscript. Near perfect reliability is achieved for mean maps with as few as 20 subjects, but variance maps require more than 1000 participants to achieve the same reliability. The same findings are found for sulcal depth mean (Figure S13C) and variance (S13D) maps as well as folding curvature mean (Figure S13E) and variance (S13F) maps.

#### Experiment I: Myelin map comparisons consistent with prior findings

Mean myelin maps from ARMS-1 (Figure S14A) and ARMS-2 (Figure S14B) show extremely high correspondence ($r^2$=0.9992). However, we could not assess subset stability nor reliability because myelin content is based on the intensity contrast ratio between the T1 and T2 image, which may vary based on platform and scanner. Because myelin content is defined as the contrast between raw intensity values from T1/T2, such unscaled values may cause problems in estimation. Therefore, future studies should examine whether researchers should perform further data harmonization, such as spatial normalization, before using myelin maps for data analysis. However, subject-specific analyses, or group-average analyses may still be appropriate(Glasser et al., 2014).

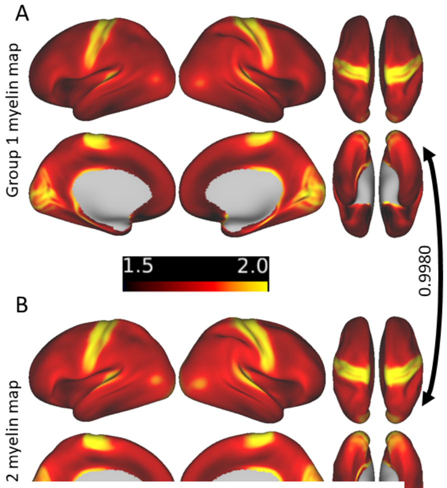


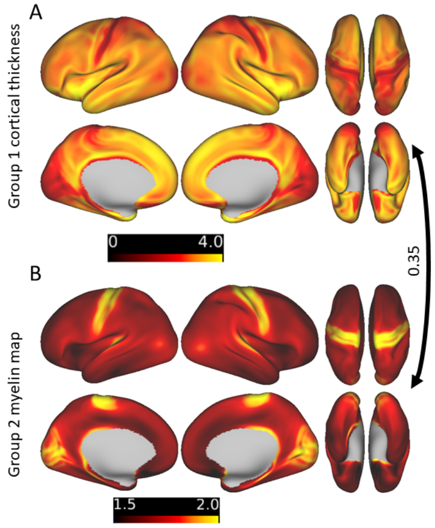


**Figure S14. Replicability of myelin maps from ARMS-1 and ARMS-2 datasets. (A)** Visualization of the mean myelin map for ARMS-1 on cortical surfaces. **(B)** The same visualization for ARMS-2. Arrow to the right of the plot denotes the proportion of variance explained between the two maps.

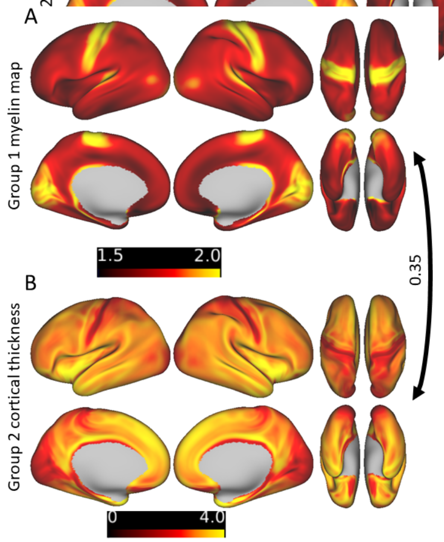


**Figure S15. Similarity between ARMS-1 mean myelin map and ARMS-2 mean cortical thickness map. (A)** Visualization of the mean myelin map for ARMS-1 on cortical surfaces. **(B)** The same visualization for the ARMS-2 mean cortical thickness map. Arrow to the right of the plot denotes the proportion of variance explained between the two maps.

**Figure S16. Similarity between ARMS-2 mean myelin map and ARMS-1 mean cortical thickness map. (A)** Visualization of the mean cortical thickness map for ARMS-1 on cortical surfaces. **(B)** The same visualization for the ARMS-2 mean myelin map. Arrow to the right of the plot denotes the proportion of variance explained between the two maps.

Additionally, we found that average myelin maps showed an inverse association with average cortical thickness. Both comparisons of the ARMS-1 mean myelin map (Figure S15A) to the ARMS-2 mean cortical thickness map (Figure S15B) and the ARMS-1 mean cortical thickness map (Figure S20A) to the ARMS-2 mean myelin map (Figure S16B) show the same degree of reproducibility ($r^2$=0.35) . As noted previously (Glasser and Van Essen, 2011; Glasser et al., 2014), this relationship may be driven by both technical and biological limitations(Glasser and Van Essen, 2011). Cortical thickness is calculated from the distance between matched vertex pairs for pial and white surfaces. Areas that have hyperintensities, such as area 6 or primary visual cortex, may underestimate both cortical thickness and myelin content (Glasser and Van Essen, 2011). To attenuate this technical limitation, the ABCD pipeline uses the same high-resolution refinement adopted previously (Glasser et al., 2013, 2014). Nevertheless, the association between cortical thickness and myelin content may remain high because myelin content and cortical thickness are biologically related. As myelin content increases, the white matter surface is likely to be defined as "closer" to the pial, leading to reductions in cortical thickness(Glasser and Van Essen, 2011).

#### Experiment II: Within study reproducibility and subset reliability of brain behavior association patterns

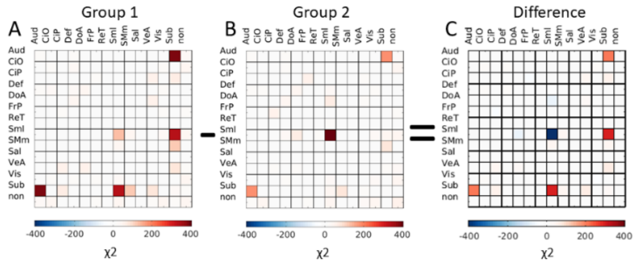


**Figure S17. Enrichment results for associations between connectivity and general ability show some consistency between ARMS-1 and ARMS-2. (A)** Matrix of significant community-by-community χ2 statistics for ARMS-1. Higher statistics indicates greater clustering for the given set of between or within-network connections. Matrices are sorted in the same order as other figures. **(B)** χ2 matrix for ARMS-2, showing overlap for within somatomotor (SmI) and between subcortical (sub) and auditory (aud),and somatomotor (SMm) communities. **(C)** Difference in chi-squared statistics, which show differences in the strength of the effect between ARMS-1 and ARMS-2.

#### Within-study reproducibility of executive function RSFC association patterns

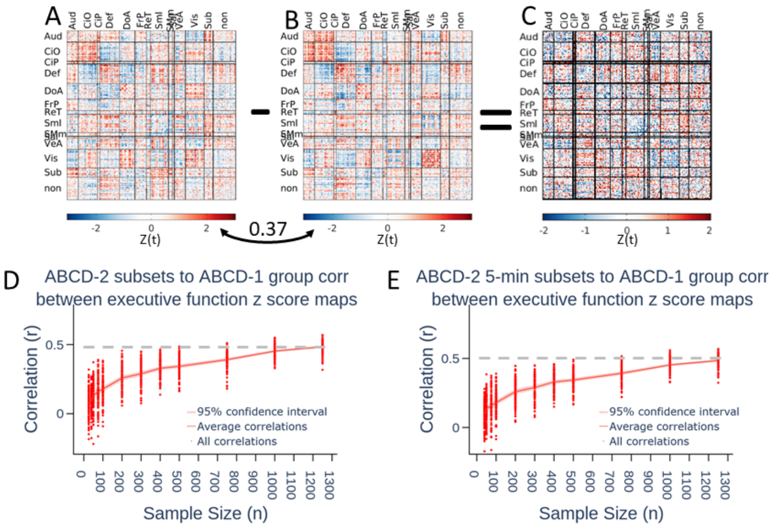


**Figure S18. Brain behavior associations between parcellated connectivity and executive function. (A)** Matrix of Z-scores for parcellated connections for ARMS-1. Matrices are sorted by community assignments. **(B)** Z-score matrix for ARMS-2. The proportion of variance explained between the two maps is indicated by the double-arrow curve. **(C)** The difference in z-scores between ARMS-1 and ARMS-2. **(D)** Plot of the correlation between subsets from ARMS-2 to the full ARMS-1 using 10 minutes of data. X-axis depicts sample size while Y shows the correlation between subsets. The grey line indicates the threshold where correlations exceed 80 percent of the correlation from the full comparison. **(E)** The same plot shown for subsets using 5 minutes of data instead of 10.

The association between ROI-based connectivity and executive function showed some within-study reproducibility ($r^2$=0.37) for ARMS-1 (Figure S18A) and ARMS-2 (Figure S18B), though less than what was found for general cognition. Similarly, differences between statistical maps were larger than found for general cognition (Figure S18C). Subset reliability analyses for both 10 minute (Figure S18D) and 5 minute (Figure S18E) ARMS-2 subsets to the 10 minute ARMS-1 subset showed similar reliability. More than 1300 participants are needed to capture 80 percent of the within-study reproducibility ($r^2$=0.3) suggesting that larger sample sizes are required to observe robustly reproducible results.

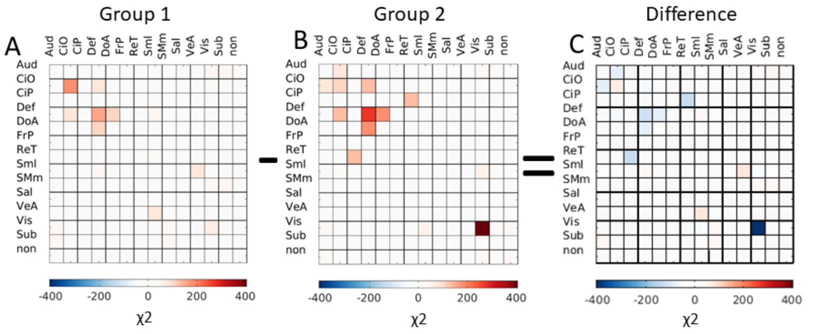


**Figure S19. Enrichment results for association between connectivity and executive function show some consistency and inconsistency between ARMS-1 and ARMS-2. (A)** Matrix of significant community-by-community χ2 statistics for ARMS-1. Higher statistics indicates greater clustering for the given set of between or within-network connections. Matrices are sorted in the same order as other figures. **(B)** χ2 matrix for ARMS-2, showing overlap for within somatomotor (Sml) and between subcortical (sub) and auditory (aud),and somatomotor (SMm) communities. **(C)** Difference in chi-squared statistics, which show differences in the strength of the effect between ARMS-1 and ARMS-2, but also different effects for connections between cingulo-opercular and auditory and between retrosplenial and default mode network.

Chi-squared results for ARMS-1 (Figure S19A) and ARMS-2 (Figure S19B) showed some consistent and inconsistent findings (Figure S19C). Notably, connections within the Cingulo-opercular (CIO) and the default mode (DMN) systems as well as between the DMN and the dorsal attention (DoA) were significantly associated with executive function. However, associations with connections sometimes appear to cluster in one system for one group (e.g. ARMS-2) and not the other (ARMS-1), such as within the visual (VIS) system.

#### Within-study reproducibility of learning/memory RSFC association patterns

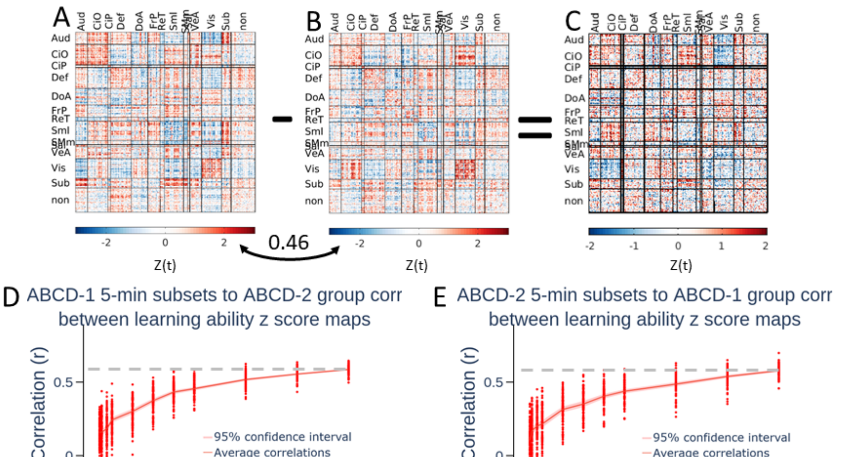


**Figure S20. Brain behavior associations between parcellated connectivity and learning/memory. (A)** Matrix of Z-scores for parcellated connections for ARMS-1. Matrices are sorted by community assignments. **(B)** Z-score matrix for ARMS-2. The proportion of variance explained between the two maps is indicated by the double-arrow curve. **(C)** The difference in z-scores between ARMS-1 and ARMS-2. **(D)** Plot of the correlation between subsets from ARMS-2 to the full ARMS-1 using 10 minutes of data. X-axis depicts sample size while Y shows the correlation between subsets. The grey line indicates the threshold where correlations exceed 80 percent of the correlation from the full comparison. **(E)** The same plot shown for subsets using 5 minutes of data instead of 10.

Linear relationships between learning/memory and ROI-based connectivity show stronger reproducibility ($r^2$=0.46) between ARMS-1 (Figure S20A) and ARMS-2 (Figure S20B). Differences between statistical maps appeared similar in degree but not always kind when compared to executive function (Figure S20C). Subset reliability analyses for both 10 minute (Figure S20D) and 5 minute (Figure S20E) ARMS-2 subsets to the 10 minute ARMS-1 subset showed similar reliability. More than 1300 participants are needed to capture 80 percent of the within-study reproducibility ($r^2$=0.37) suggesting that larger sample sizes are required to observe robustly reproducible results.

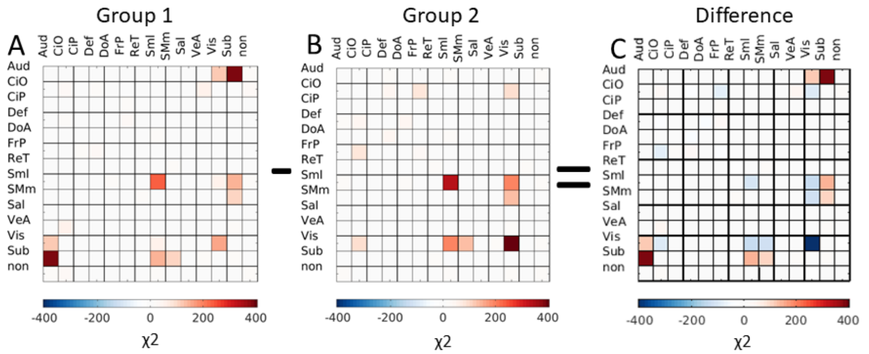


**Figure S21. Enrichment results for association between connectivity and learning/memory show some consistency and inconsistency between ARMS-1 and ARMS-2. (A)** Matrix of significant community-by-community χ2 statistics for ARMS-1. Higher statistics indicates greater clustering for the given set of between or within-network connections. Matrices are sorted in the same order as other figures. **(B)** χ2 matrix for ARMS-2, showing overlap for within somatomotor (SmI) and between subcortical (sub) and auditory (aud), and somatomotor (SMm) communities. **(C)** Difference in chi-squared statistics, which show differences in the strength of the effect between ARMS-1 and ARMS-2, but also different effects for connections between visual, subcortical, somatomotor, and auditory systems.

Chi-squared results for ARMS-1 (Figure S21A) and ARMS-2 (Figure S21B) showed some similarities. Notably, within SMI and within Vis systems showed similar effects for both ARMS-1 and ARMS-2. However, many associations were divergent between ARMS-1 and ARMS-2. ARMS-1 showed between-system associations clustering for Aud-vis, sub-SMI, and sub-SMm systems. ARMS-2 showed between-system associations clustering for Vis-CIO, VIS-SMI, and VIS-SMm systems.

#### Subset reliability analyses for ARMS-1 subsets to ARMS-2

The manuscript so far has only discussed ARMS-2 subset to ARMS-1 reliability analyses to make the manuscript easier to read. Some may be concerned if ARMS-1 subsets to ARMS-2 show different degrees of reliability. Here, ARMS-1 subset to ARMS-2 reliability analyses are shown to illustrate that the findings are the same and alleviate this concern (Figure S22). Subset reliability analyses at 10 (Figure S22A) and 5 (Figure S22B) for the association between general cognition and ROI-based connectivity replicate the findings in the main manuscript. With 10 minute of data, 80 percent of subset reliability (Figure S22A; dotted grey line) is captured at 1250 participants, but not with 5 minutes of data (Figure S22B; dotted grey line). For associations between executive function and ROI-based connectivity, both 10 (Figure S22C) and 5 (Figure S22D) minutes of data need more than 1300 participants to capture more than 80 percent of subset reliability (Figures S12C and D; dotted grey lines). The same findings are shown for associations between learning/memory and ROI-based connectivity for 10 (Figure S22E) and 5 (Figure S22F) minutes of data.

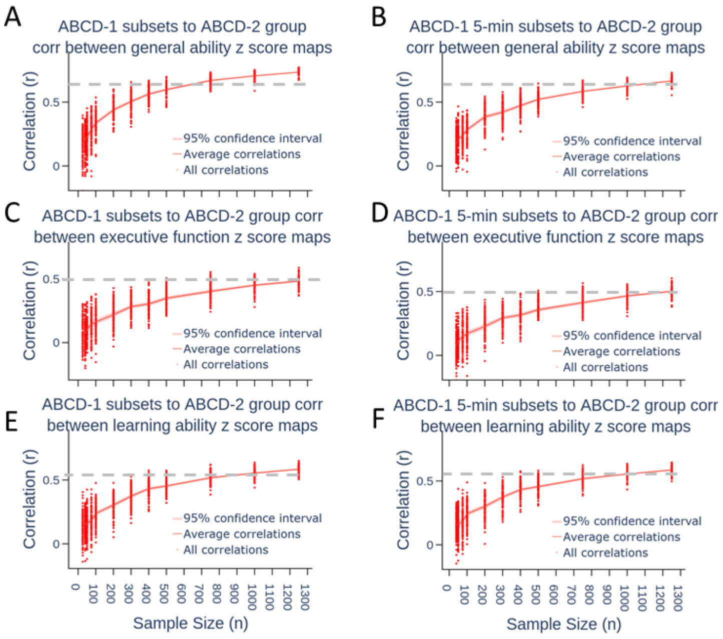


**Figure S22. Remaining subset analyses replicate findings from ARMS-1 subsets to the full ARMS-2 dataset.** All figures depict the correlation from ARMS-1 subsets to ARMS-2 dataset. The x-axis represent the given subset’s sample size, while the y-axis represents the correlation between the given subset an the 10 minute ARMS-2 dataset. Grey dashed lines indicate 80 percent of the correlation between the statistical maps for the 10 minute full ARMS-1 and ARMS-2 datasets. **(A)** Replicability for general ability with 10 minute subsets. **(B)** Replicability for general ability with 5 minute subsets. **(C)** Replicability for executive function with 10 minute subsets. **(D)** Replicability for executive function with 5 minute subsets.**(E)** Replicability for learning/memory with 10 minute subsets. **(F)** Replicability for learning/memory with 5 minute subsets.

AND ATTENTION. Monogr. Soc. Res. Child Dev. *78*, 16–33.
